# Disruption of Pre-Bötzinger Complex neuropeptidergic tonality controls fear and metabolic response

**DOI:** 10.64898/2026.01.01.697304

**Authors:** Sanutha Shetty, Amir Farmanbar, Pamela Toh, Ramazan Yildiz, YoungUK Jang, Samuel Duesman, Aidan Warnock, Diego Espinoza, Sarah A. Stanley, Prashant Rajbhandari, Abha K. Rajbhandari

## Abstract

Stress profoundly impacts systemic metabolism, yet the central circuits linking stress responses to peripheral metabolic regulation remain poorly defined. Here, we identify the preBötzinger complex (preBötC), a brainstem breathing rhythm generator, as a key stress-responsive hub coordinating metabolic adaptations. Using viral tracing, we show that preBötC neurons project to brown adipose tissue and liver, and that a subset of these projection neurons expresses the pituitary adenylate cyclase–activating polypeptide (PACAP) receptor PAC1R, positioning PACAP signaling as a critical modulator of this circuit. Whole-brain c-Fos mapping revealed robust preBötC activation under stress, while spatial transcriptomics demonstrated altered neuronal metabolic circuitry in preBötC following PAC1R ablation. PAC1R knockdown in preBötC combined with stress resulted in blunted respiratory rhythmicity, reduced sympathetic innervation, and suppression of energy expenditure and lipid metabolic pathways in brown fat, while reprogramming hepatic transcriptional networks toward amino acid metabolism and gluconeogenesis. These findings define a unique neuropeptidergic brainstem–periphery circuit integrating stress, respiration, and metabolism.

**Highlights:**

- PAC1R deletion in preBötC amplifies PTSD-like fear—greater generalization and freezing despite equal stress.
- PreBötC-PAC1R neurons send projections to BAT and liver, forming a respiratory-metabolic hub.
- Loss of PAC1R lifts the preBötC “brake,” raising resting breathing rate and magnifying stress-induced heart-rate spikes.
- Loss of PAC1R show systemic metabolic failures—glucose intolerance, lower VO₂ input, reduced energy expenditure, and weaker BAT sympathetic tone.
- Spatial transcriptomics reveal marked shifts in preBötC neuronal subpopulations after PAC1R ablation.

## Introduction

Breathing is a fundamental physiological process that allows us to take in oxygen to fuel our cells and eliminate carbon dioxide as a byproduct of cellular metabolism (Feldman and Smith, 1989). Controlled by the autonomic nervous system (ANS), breathing is automatic and unconscious, even when we are asleep and resting. While requiring no conscious effort, the respiratory rhythm is highly dynamic and can be modified by emotional states (Klein, 1993; Ley, 1988), cognitive needs (Heck et al., 2019; Melnychuk et al., 2018; Zelano et al., 2016), and physiological demands (Borgdorff, 1975; Dominelli and Sheel, 2024).

Disruptions in breathing rhythms are a hallmark feature of most psychiatric and stress-related disorders (Blazer and Hybels, 2010; van Liempt et al., 2011; von Leupoldt et al., 2011; Von Leupoldt et al., 2010). Among these, post-traumatic stress disorder (PTSD) is a disorder characterized by augmented arousal (First, 2013; Litz et al., 1996; Shepherd and Wild, 2014), dysregulated stress responses (Yehuda, 2002), abnormal metabolic regulation (Michopoulos et al., 2016; Rosenbaum et al., 2015) , and hampered breathing—rapid and/or shallow—during episodes of flashbacks, panic, or anxiety (Blechert et al., 2007; Tjondrorahardja et al., 2024; van Liempt et al., 2011). Moreover, there is evidence suggesting that intentionally slowing the breathing rate can enhance positive affect regulation in response to stress (Brown and Gerbarg, 2005; Brown et al., 2013). Many yogic practices are based on this principle and are gaining wide support for their use in clinical applications for stress, anxiety, and cardiometabolic disorders (Hamasaki, 2020). These findings show a bidirectional relationship between stress reactivity and breathing. However, the mechanisms by which respiratory dynamics regulate stress-related behavior and metabolism are unknown despite these results. This is especially important in trauma-related disorders like PTSD, where stress sensitivity and metabolic dysregulation are usually correlated and result in comorbidities like cardiovascular disease, diabetes, and obesity (Michopoulos et al., 2016; Rosenbaum et al., 2015). Building on this understanding, we propose that breathing serves as a dynamic interface between stress reactivity and metabolic regulation.

The preBötzinger Complex (preBötC) in the ventrolateral medulla is considered the main site for respiratory rhythmogenesis (Feldman and Del Negro, 2006; Feldman et al., 2013; Gray et al., 2001), with sufficient evidence for an analogous role in humans as well (Schwarzacher et al., 2011; Smith et al., 1991). This structure is characterized by conspicuous cellular heterogeneity, including both inhibitory and excitatory types of neurons. These neurons express a diverse array of neuropeptidergic receptors, including the G protein–coupled PAC1R receptor (Adcyap1r1), which specifically binds to pituitary adenylate cyclase-activating polypeptide (PACAP; Adcyap1) (Shi et al., 2021). Most notably, Shi et al. (Shi et al., 2021) also demonstrated that PAC1R-expressing neurons in the preBötC are essential for initiating and maintaining stable breathing immediately after birth in mice, as they mediate the stimulatory effects of PACAP released from retrotrapezoid nucleus (RTN) neurons. As our work and others’ have demonstrated, PACAP/PAC1R signaling plays a central role in modulating stress response, energy homeostasis, and respiration (Bozadjieva-Kramer et al., 2021; Clancy et al., 2023; Duesman et al., 2022; Rajbhandari et al., 2021; Ressler et al., 2011; Shi et al., 2021). Here, we have report that PAC1R signaling within the preBötC serves as a key integrative mechanism linking respiratory rhythm with stress adaptation and metabolic regulation.

## Results

### Determining the specific pathway of respiratory rhythm-generating neurons in the preBötC that are activated during stress

With the renewed interest in the James-Lange theory as a framework for understanding the origins of emotion, the basic physiological act of breathing is increasingly recognized as a potential driver of fear expression (Fehr and Stern, 1970; James-Lange, 1922). A recent study demonstrated that episodic slow breathing, induced by photostimulating GlyT2-expressing neurons in the preBötC, significantly attenuates fear responses (de Sousa Abreu et al., 2024). Therefore, to assess whether preBötC could be mediating stressor response, we followed the experimental design as laid out in Fig. 1A. Stress Enhanced Fear Learning (SEFL) recapitulates critical aspects of PTSD-like fear in rodents, including long-term sensitization of fear learning caused by an acute stressor (Rajbhandari et al., 2018). Ninety minutes after Day 3 of the SEFL paradigm, animals were euthanized, and brains were collected. A subset of the brain sample underwent the Fast 3D clearing protocol followed by whole-brain immunostaining to detect cFos+ cells, an immediate early gene used as a marker of neuronal activity (Supplemental Fig. 1A and 1B). Using an unbiased whole-brain screening approach, we observed selective activation of preBötC neurons in SEFL-exposed mice, but not in No-SEFL controls. To investigate the role of PAC1R-expressing neurons in stress responses, we performed RNAscope fluorescent in situ hybridization (FISH) combined with immunohistochemistry (IHC) on cryosectioned brain samples containing the preBötC. Fig. 1B represents the PAC1R mRNA fluorescence and immunohistochemical staining of cFos, in sections containing the preBötC in the No-SEFL controls and the SEFL group. The quantification of cFos+ neurons in the preBötC showed a significantly higher expression of cFos in the SEFL mice than the No-SEFL mice (n=5-6; unpaired t-test, t=4.345, p=0.002) (Fig. 1C). There is also a positive correlation between preBötC cFos and % freezing in mice (n=5-6; r² = 0.68, p=0.002). Quantification of PAC1R mRNA and cFos expression in the preBötC revealed that almost two-thirds of stress-activated (cFos+) neurons express PAC1R mRNA (Fig. 1D). The PACAP/PAC1R receptor pathway plays a role in both human and rodent stress responses alike (Hammack et al., 2009; Mustafa, 2013; Ressler et al., 2011). To identify PACAPergic inputs to preBötC-PAC1R-expressing neurons, we utilized a retrograde tracing approach restricted to a single synapse. This strategy allowed us to map monosynaptic inputs to the preBötC using CTB-A647 in ADCYAP1-GFP mice, as illustrated in the schematic (Fig. 1E). CTB virus injection in ADCYAP1-GFP mouse brains confirmed viral labeling of neurons in the preBötC (Fig. 1F). Among the many brain regions previously shown by Yang et al. (Yang et al., 2020) to project monosynaptically to the preBötC, we identified the anterior nucleus of the solitary tract (NTS) as a prominent source of PACAPergic input, exhibiting strong colocalization of PACAP with CTB-A647 (Fig. 1G). While other regions, including the medial NTS (m-NTS), periaqueductal gray (PAG), hypothalamus, and medial preoptic area (MPO), also send monosynaptic projections to the preBötC but showed minimal PACAP colocalization (Supplemental Fig. 1C-1F). The preBötC consists of a heterogeneous population of excitatory (glutamatergic) and inhibitory (GABAergic) neurons (Feldman et al., 2013). To identify whether PAC1R receptors in the preBötC are expressed on glutamatergic or GABAergic neurons, we conducted RNAscope combined with immunohistochemistry (IHC) on tissue sections encompassing the preBötC, as illustrated by the RNAscope staining in the corresponding coronal section (Fig. 1H). RNAscope was used to label transcripts of PAC1R mRNA, and IHC was used to label either GAD2 (GABAergic) or Vglut2 (glutamatergic) in subsequent sections. We found higher colocalization of PAC1R mRNA with GAD2 (Fig. 1I) than with Vglut2 (Fig. 1J), as shown by the quantification of the proportion of PAC1R+/GAD2+v. PAC1R+/Vglut2+ neurons (n=4, unpaired t-test, t=14.33, p<0.0001) (Fig. 1K).

**Figure 1.**
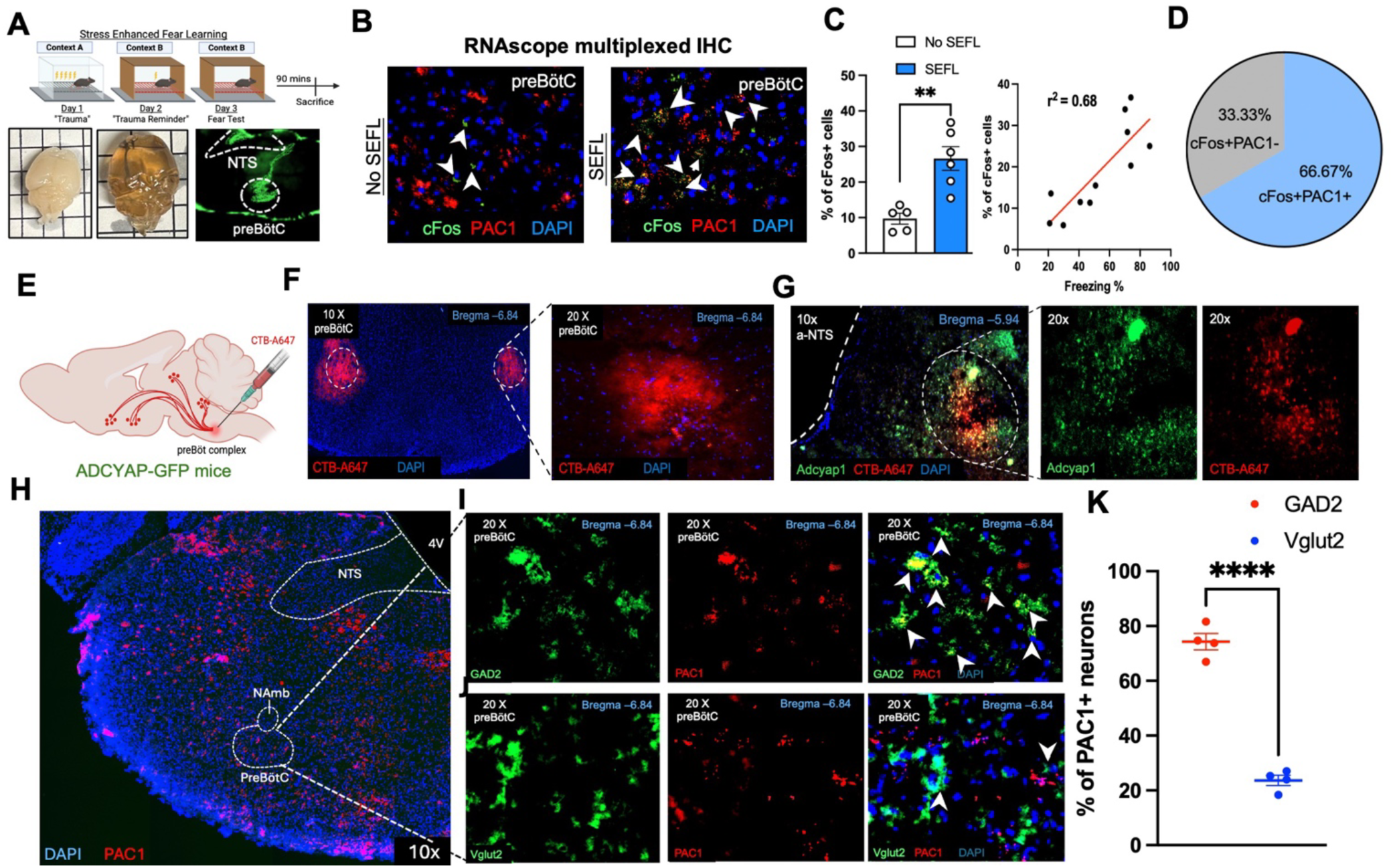
Stressors activate preBötC neurons and receive PACAPergic inputs from the NTS. (A) Experimental design for assessing stress-induced activation of PAC1R neurons in the preBötC using the SEFL paradigm; Representative images of brain before and after tissue clearing, with corresponding 3D reconstructions of cFos immunolabeling in No SEFL and SEFL groups. B) Representative RNAscope FISH multiplexed with IHC images showing PAC1R mRNA (red), cFos protein (green) and DAPI (blue) expression in the preBötC of No-SEFL and SEFL mice. C) Quantification of cFos+ neurons in the preBötC in No-SEFL mice compared to SEFL mice (n=5–6; unpaired t-test, t=4.345, p=0.002); Correlation analysis showing a positive relationship between the number of cFos+ neurons in the preBötC and percent freezing behavior in SEFL mice (n=5–6; r² = 0.68, p=0.002). D) Pie chart illustrating the proportions of cFos+ and PAC1R+ neurons, and cFos+ and PAC1R neurons. E) Schematic of retrograde tracing strategy using CTB-Alexa 647 (CTB-A647) in ADCYAP1-GFP mice. F) Representative image of viral labeling of preBötC neurons by CTB-A647 in ADCYAP1-GFP mice. G) Representative image of the section containing a-NTS expressing PACAP (green) and CTB-A647 (red). H) Representative image of a coronal brain section containing the preBötC showing RNAscope labeling of PAC1R mRNA. I–J) Dual labeling of PAC1R mRNA with GAD2 (I) or Vglut2 (J) using RNAscope multiplexed with IHC. K) Quantification of PAC1R+/GAD2+ versus PAC1R+/Vglut2+ neurons in the preBötC (n=4, unpaired t-test, t=14.33, p<0.0001).

### Ablation of preBötC PAC1R receptors disrupts stress responses and cardiorespiratory function

To assess the role of PAC1R receptors in the preBötC in regulating fear behavior, we used a Cre-loxP strategy in PAC1R^flox/flox^ mice. Animals received bilateral injections of either AAV2-hSyn-GFP or AAV2-hSyn-GFP-Cre into the preBötC (LM: ±1.25; AP: −6.78; DV: −5.70 mm from Bregma; Fig. 2A). To assess PTSD-like stress responses, we employed the SEFL model, which allowed us to evaluate alterations in fear reactivity by measuring the percentage of freezing and shock reactivity. We also utilized a pulse oximeter in freely behaving animals during the SEFL paradigm to monitor cardiorespiratory changes associated with behavioral responses. We assessed anxiety-like phenotype in these mice by performing a battery of assays, including the open field light-gradient task (Nagai et al., 2019) and the elevated plus maze (EPM) task. Additionally, we analyzed metabolic outcomes through a combination of indirect calorimetry, glucose tolerance tests to assess metabolic flexibility, pyruvate tolerance test, insulin tolerance test, and quantified levels of mouse insulin in serum through ELISA to characterize the metabolic profile of these mice. Three weeks after bilateral virus injections, PAC1R expression was observably reduced in the preBötC neurons of Cre animals. (Fig. 2B). Further, quantification of the mean fluorescence intensity (MFI) of PAC1R mRNA visualized by RNAscope confirmed a significant reduction in PAC1R mRNA signal in Cre-injected mice compared to controls injected with control GFP virus (n=4; unpaired t-test, t=5.216, p= 0.0022) (Fig. 2C). In the SEFL paradigm, freezing behavior on Day 1 did not differ between groups (two-way ANOVA, F(1,360)=8.495, p=0.38) (Supplemental Fig. 2A). However, we did observe a significant group effect on shock reactivity on Day 1 (n=18-20, two-way ANOVA, F(1,360)=34.17, p<0.0001) (Supplemental Fig. 2B). *Posthoc* analysis revealed a significant reduction in shock reactivity in the Cre-SEFL group on the 10^th^ footshock in comparison to Ctrl-SEFL group (n=18-20, Tukey’s multiple comparison test, p=0.0167). On Day 2, freezing in a novel context (context B) revealed a main group effect (n=12-20, two-way ANOVA, F(3,51)=17.46, p<0.0001); Cre-SEFL animals (both males and females) froze significantly more than Cre-No SEFL controls (n=6-13, Tukey’s multiple comparison test, p<0.0001), indicating increased fear generalization (Fig. 2D). No group differences in shock reactivity were observed on Day 2 (n=12-20, F(3,52)=0.6125, p=0.604; Fig. 2E). The key assessment in the SEFL model is the context test on Day 3. During this test, we observed a main effect of stress and PAC1R deletion (n=12-20, two-way ANOVA, F(3,52)=29, p<0.0001) (Fig. 2F). *Posthoc* comparison showed that the Ctrl-SEFL animals exhibited significantly higher freezing behavior compared to Ctrl-No SEFL controls (n=13-15, Tukey’s multiple comparison test, p<0.0001). Further, *posthoc* comparisons showed a significant enhancement in freezing percentage in Cre-No SEFL mice compared to the Ctrl-No SEFL group (n=13-15, Tukey’s multiple comparison test, p=0.014[females]; p=0.0009[males]), suggesting that PAC1R loss sensitizes animals to even mild stress. We observed no statistically significant *posthoc* difference in freezing behavior between Ctrl-SEFL or Cre-SEFL (n=15-20, Tukey’s multiple comparison test, p=0.848), potentially due to a ceiling effect. Therefore, to better assess the effects of PAC1R deletion on freezing behavior, we employed a modified SEFL protocol with a lower-intensity stressor on Day 1, in which mice received five 1 mA footshocks over 30 minutes. We performed the day 2 and 3 tests as usual (Supplemental Fig. 2D). On Day 2, we recaptured similar effects as our original experiment, wherein we observed Cre-SEFL animals showed greater freezing compared to the Ctrl-SEFL group, indicating higher fear generalization in the Cre group (n=5; unpaired t-test, t=6.135, p=0.0009) (Supplemental Fig. 2E). We observe no difference in shock reactivity (Supplemental Fig. 2F, n=5, unpaired t-test, t=1.72, p=0.136). However, on Day 3, we now observe significantly increased freezing levels in the Cre-SEFL group compared to Ctrl-SEFL animals, confirming that PAC1R deletion enhances fear expression even under lower stress conditions (n=5; unpaired t-test, t=2.73, p=0.034) (Supplemental Fig. 2G). We also delineated any differences in fear acquisition after SEFL in a subset of Ctrl-SEFL and Cre-SEFL animals by placing them back in context A for 5 minutes and observed no significant group effects (n=5, unpaired t-test, t=0.086, p=0.933) (Fig. 2G). To assess the cardiac response to the SEFL, we measured heart rate in behaving mice during SEFL Day 3 to correlate freezing behavior with cardiac output. We identified significantly higher ΔHR in the Cre-SEFL group compared to the Ctrl-SEFL group during phases of immobility in the mice. Fig. 2H represents an example trace of ⊗HR in Cre-SEFL and Ctrl-SEFL mice, and Fig. 2I shows the average ⊗HR that reveals a main effect of PAC1R deletion (n=12-13, one-way ANOVA, F(3,40)=4.593, p=0.07). *Posthoc* comparisons also showed this increased ⊗HR to be statistically significant between Cre-SEFL and Ctrl-No SEFL groups as well (n=10-13, Tukey’s multiple comparison test, p=0.025) (Supplemental Fig. 2C). We observed a similar increase in ⊗HR in the Cre-SEFL group compared to the Ctrl-SEFL in the modified SEFL protocol (5 footshocks) (n=5, unpaired t-test, t=2.97, p= 0.03) (Supplemental Fig. 2H). We also measured the breathing rate in Ctrl-SEFL and Cre-SEFL mice under anesthesia using the pulse oximeter system. Representative breathing trace over 5 minutes revealed an increased breathing rate per minute (BRPM) in Cre-SEFL mice compared to Ctrl-SEFL mice (Fig. 2J). The average BRPM over 5 minutes showed a statistically significant difference between the Cre-SEFL and Ctrl-SEFL groups (n=6–8, unpaired t-test, t=2.625, p=0.02) (Fig. 2K). We observed no effect of stress or PAC1R deletion in anxiety-like phenotype in any of the groups in both open-field light gradient tasks (n=9-19, two-way ANOVA, F(3,153)= 0.556, p= 0.644; F(3,153)= 0.569, p= 0.635) (Supplemental Fig. 2I-K) or EPM (n=7-14, one-way ANOVA, F(3,30)= 1.15, p= 0.354; F(3,31)= 0.257, p= 0.855) (Supplemental Fig. 2L-N).

**Figure 2.**
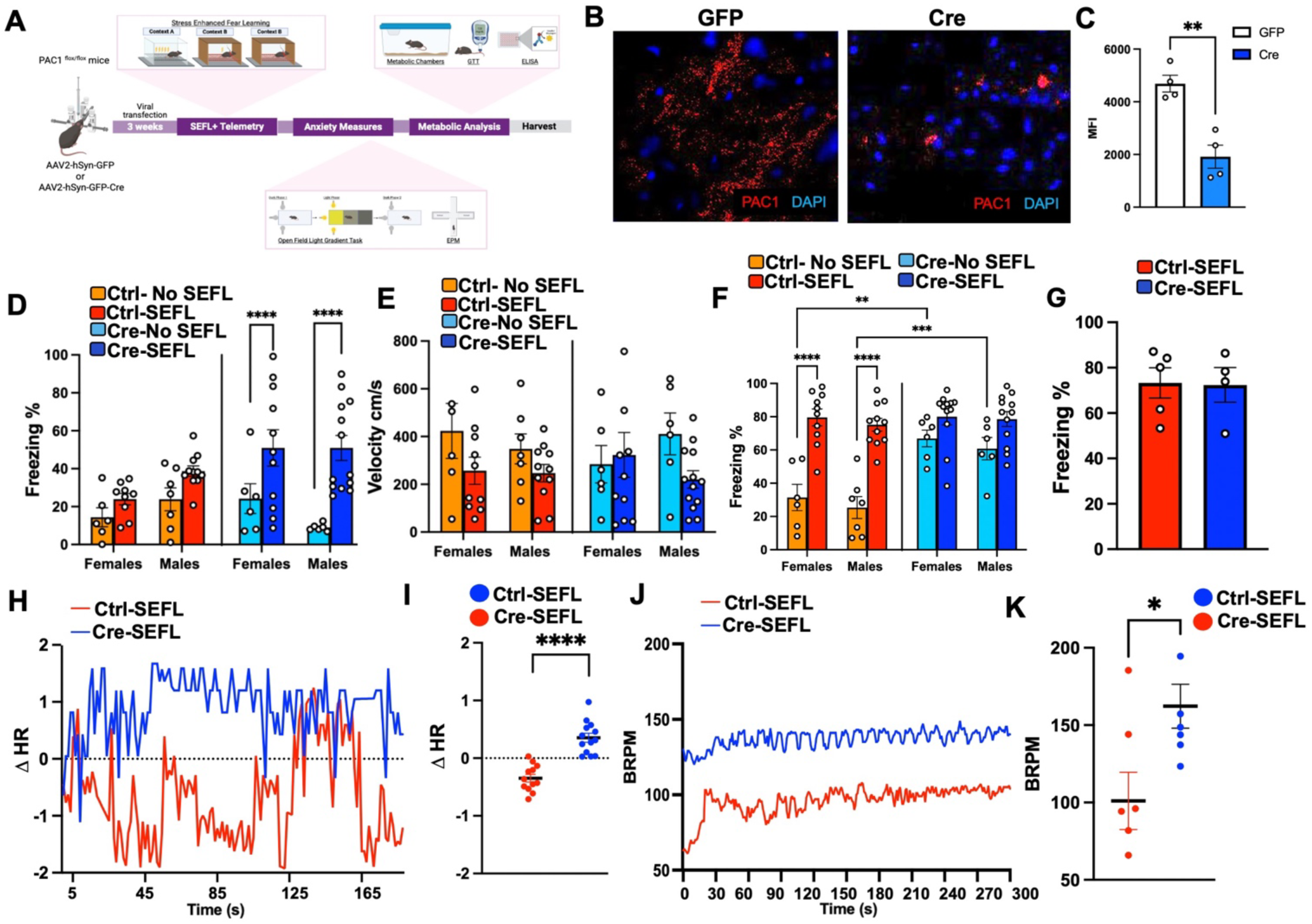
PAC1R deletion in the preBötC exacerbates defensive fear behavior and physiological responses. A) Experimental timeline of the loss-of-function study assessing fear behavior, anxiety-like phenotype, cardiorespiratory function, and metabolic adaptations. B) Representative images of PAC1R mRNA expression in mice receiving rAAV2-hSyn-GFP-Cre or rAAV2-hSyn-GFP injections (Ctrl) into the preBötC. C) Quantification of mean fluorescence intensity (MFI) of PAC1R mRNA in both Cre and Ctrl groups (n=4; unpaired t-test, t=5.216, p= 0.0022). D) Percent freezing measured on Day 2 of SEFL in male and female mice (n=12-20, two-way ANOVA, F(3,51)=17.46, p<0.0001). E) Shock reactivity assessed by measuring velocity in both male and female mice to evaluate their immediate response to the footshock stimulus on day 2 SEFL (n=6-10; two-way ANOVA, F(3,52)=0.6125, p=0.604). F) Percent freezing on Day 3 of SEFL in male and female mice (n=12-20, two-way ANOVA, F(3,52)=29, p<0.0001). G) Fear acquisition in the training context (Context A) was not affected by deletion (n=5; unpaired t-test, t=0.086, p=0.933). H) Example trace of deviation in heart rate (ΔHR) during 8s freezing bouts in Ctrl-SEFL and Cre-SEFL animals. I) Average ΔHR during the entire duration of freezing on Day 3 of the SEFL paradigm (n=12-13, one-way ANOVA, F(3,40)= 4.593, p=0.07). J) Example trace of breathing rate per minute (BRPM) measured in anesthetized Ctrl-SEFL and Cre-SEFL mice for 5 minutes. K) Average BRPM during 5 minutes of recording in Ctrl-SEFL and Cre-SEFL groups (n=6–8, unpaired t-test, t=2.625, p=0.02).

### Spatial transcriptomics reveals a major metabolic remodeling of the preBötC complex neuronal signaling upon PAC1R signaling under stress

Spatial transcriptomic profiling of the preBötC following PAC1R deletion after stress (Cre-SEFL) (Fig. 3A and 3B and Supplemental Fig. 3A-3H) revealed profound changes in abundance of neuronal population and genes regulating both neuronal signaling and metabolism (Fig. 3C and 3D). Among other, Cre-SEFL preBötC showed prominent changes in cholinergic, glutamergic, and GABAergic neuronal populations (Fig. 3D). To examine the transcription program controlled by PAC1R signaling in preBötC, we performed differential gene expression analysis (DEG) on all preBötC clusters, cholinergic, glutamatergic, and GABAergic clusters, as represented in the volcano plots (Fig. 3E-3G). A striking feature of the all preBötC clusters volcano plot was the upregulation of multiple mitochondrial-encoded genes (mt-Co2, mt-Cytb, mt-Co3, mt-Nd1, mt-Nd2, mt-Atp6) (Fig. 3E). This suggests heightened oxidative phosphorylation and bioenergetic activity within preBötC neurons. Given that preBöC neurons are rhythmically active and require continuous ATP for ion channel function and neurotransmitter release, this shift may reflect a compensatory metabolic adaptation to PAC1R loss. Supporting this, other upregulated genes such as App, C1ql1, and Slc12a are linked to synaptic regulation, glutamatergic transmission, and ion homeostasis—all processes critical for rhythm generation. Genes suppressed in PAC1R-deficient preBötC included Dlk1, Ndn, Nap1l5, Gnas, and Adora2a, which are known regulators of neuronal differentiation, plasticity, and GPCR signaling. Their downregulation points toward reduced neuroplastic capacity and impaired neuromodulatory responsiveness. Pathway enrichment revealed suppression of glycogen metabolism, lipid droplet organization, and glucose utilization, suggesting impaired energetic adaptability at the circuit level. For example, Adora2a encodes the adenosine A2A receptor, a modulator of excitability in respiratory networks, while Gnas regulates cAMP signaling, which is central to fine-tuning rhythm output. The preBötC is uniquely sensitive to neuromodulatory peptides, including PACAP (which acts via PAC1R), to stabilize breathing under stress. The observed transcriptomic shift enhanced mitochondrial activity but dampened signaling/plasticity pathways, suggesting that in the absence of PAC1R input, neurons attempt to sustain activity through metabolic upregulation, but at the expense of adaptability and neuromodulatory control. This imbalance may underlie the breathing instability and stress hypersensitivity observed with PAC1R deletion. Comparative transcriptomic profiling of cholinergic-enriched preBötC neurons also revealed distinct alterations with PAC1R deletion (Fig. 3F). Several synaptic and neurodevelopmental regulators were significantly downregulated, including Prph (peripherin), Slc6a7 (a glutamate transporter), Syp (synaptophysin), Nsg1, Rin1, and Nell2, all of which contribute to synaptic vesicle cycling, axonal integrity, and plasticity. Importantly, the reduction of Prph, a cytoskeletal protein required for axonal stability in cholinergic neurons, and Syp, a synaptic vesicle marker, points to impaired structural and functional connectivity within the respiratory network. Enrichment analysis highlighted suppression of glycogen breakdown, ROS regulation, and nitric oxide signaling, pointing to a reduced capacity for sustaining bioenergetically demanding cholinergic drive. Together, these changes highlight that PAC1R loss in cholinergic neurons drives a metabolic–synaptic tradeoff: neurons increase mitochondrial capacity but lose synaptic adaptability and axonal support. In the context of the preBötzinger complex, where precise cholinergic modulation is essential for stabilizing inspiratory rhythm, this imbalance likely weakens the flexibility and resilience of the respiratory oscillator, making it more susceptible to stress-induced dysfunction. Within GABAergic neurons, genes such as Dlk1, Kcnk9, Gch1, and Syt4 were significantly repressed (Fig. 3G). These transcripts converge on ion channel regulation, mitochondrial coupling, and redox balance. Pathway enrichment identified suppression of neuronal excitability, resting membrane potential regulation, and carbohydrate metabolism, reflecting compromised inhibitory tone and metabolic stability. Glutamatergic neurons displayed a distinct compensatory transcriptional profile, with upregulation of Abhd17a, Oxct1, Pde4b, Trpc3, Itgb8, and Neto2 (Fig. 3H). These genes are linked to cAMP homeostasis, ketone body utilization, calcium signaling, and axonal remodeling. Pathway analysis revealed enhanced Ras signaling, receptor trafficking, and synaptic plasticity, suggesting an adaptive excitatory rewiring to counterbalance metabolic deficits in inhibitory and cholinergic neurons. Together, these findings show that PAC1R acts as a metabolic stabilizer in the preBötC, sustaining neuronal fuel utilization and excitability across subtypes. Its loss induces a coordinated loss of control of inhibitory/cholinergic metabolic tone while promoting adaptive excitatory plasticity.

**Figure 3.**
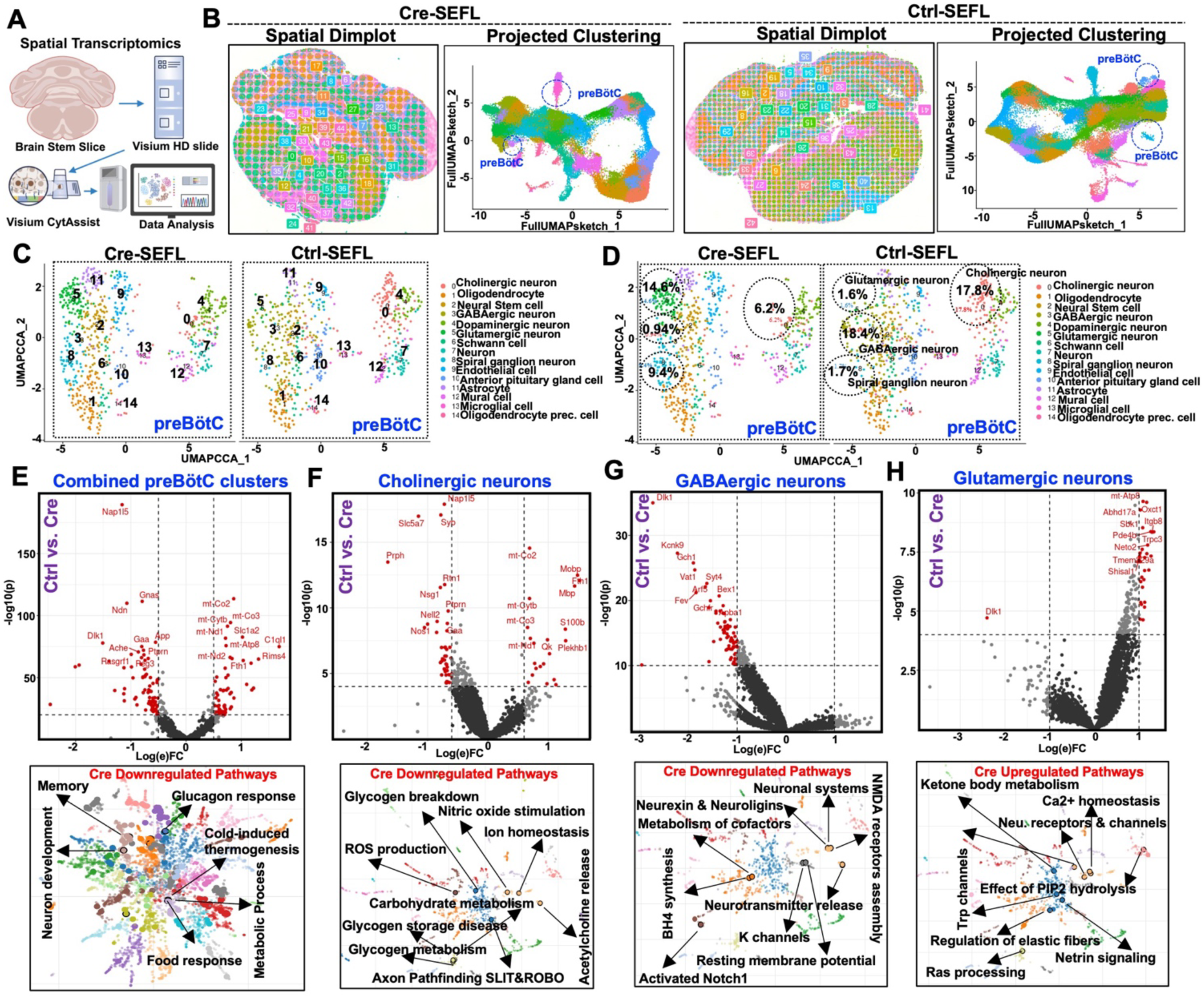
Single-cell transcriptomic mapping of preBötC neuronal populations reveals cell-type–specific transcriptional reprogramming upon PAC1R deletion. A) Schematic of Visium HD spatial transcriptomics workflow. B) Spatial dimplot and projected clustering of brain regions with UMAP visualizations of preBötC populations. PreBötC clusters are circled and annotated across datasets. C) UMAP showing distinct clustering of cholinergic, GABAergic, and other cell types subtypes in the isolated preBötC. D) UMAP showing changes in percentage of the circled clusters in Cre-SEFL and Ctrl-SEFL preBötC. E–H) Volcano plots displaying differential gene expression (top) and pathway enrichment analysis in combined preBötC clusters (E), cholinergic neurons (F), GABAergic neurons (G), and glutamatergic neurons (I). Red points represent significantly altered genes (adjusted p < 0.05).

### PAC1R ablation in the preBötC paired with SEFL disrupts glucose and energy metabolism

Our spatial transcriptomic analysis of the preBötC following PAC1R deletion revealed selective downregulation of genes critical for cholinergic signaling, GABAergic modulators, and synaptic vesicle machinery, alongside upregulation of genes in glutamatergic neurons. Functionally, these transcriptional shifts point to a rebalancing of excitatory and inhibitory tone within the preBötC, with metabolic consequences. Recent research has mapped the high-resolution connectivity of efferent projections to the preBötC (Yang and Feldman, 2018) and afferent inputs to other brainstem regions involved in respiratory control (Tan et al., 2010) within the central nervous system. However, fully understanding how the brain regulates breathing and integrates it with whole-body metabolism requires expanding our focus beyond these neural circuits to include peripheral connectivity with metabolic tissues—a critical yet largely unexplored aspect of respiratory control of metabolic regulation. Building upon our spatial transcriptomics to assess whether perturbation of preBötC controls peripheral metabolism, we employed a multi-synaptic retrograde tracing approach using pseudo-rabies virus (PRV) to label distant cell bodies projecting from metabolic tissues, including the brown adipose tissue (BAT), liver and inguinal white adipose tissue (iWAT) (Fig. 4A). Retrograde labeling of the preBötC neurons was evident following PRV-RFP injection into the BAT with a high degree of colocalization with PAC1R mRNA (Fig. 4B and 4E). Further, PRV-GFP injection into the larger lobe of the liver also resulted in GFP labeling of preBötC neurons and a high degree of colocalization with PAC1R mRNA (Fig. 4C and 4F). PRV-GFP injection into white adipose tissue (WAT) did not result in much GFP labelling of preBötC neurons (Fig. 4D). To examine the consequence of Cre-SELF on BAT metabolism, we performed RNA-seq on BAT from Ctrl-SEFL and Cre-SEFL mice. We identified 1,254 differentially expressed genes (DEGs; FDR < 0.05) in Ctrl-SEFL versus Cre-SELF mice. We focused on genes that were highly downregulated in Cre-SELF compared to control. Gene-set enrichment analyses highlighted concerted down-regulation of PPAR signaling, fatty-acid β-oxidation, and AMPK-mediated lipid-droplet organization and gene associated with BAT thermogenesis and metabolism (Fig. 4G, 4H and Supplemental Fig. 4A and 4B). Next, to assess the impact of PAC1R deletion combined with SEFL on efferent sympathetic innervation, we collected BAT from Ctrl-SEFL and Cre-SEFL mice, performed tissue clearing, and stained for tyrosine hydroxylase (TH), a marker of sympathetic nerve fibers (Supplemental Fig. 4C). Fig. 3F shows representative images of BAT from both groups labeled for TH. Quantification of TH signal intensity, normalized to tissue volume, revealed markedly reduced TH expression in the BAT of Cre-SEFL mice compared to GFP-SEFL controls (n=2, unpaired t-test, t=19.92, p<0.0001) (Fig. 4I and 4J). The levels of corticosterone were similar between Cre-SEFL and Ctrl-SEFL (Supplemental Fig. 4D). To further test the consequence of diminished BAT function on energy homeostasis, we performed indirect calorimetry using metabolic chambers. We observed a significant effect of PAC1R deletion between Ctrl-SEFL and Cre-SEFL mice on VO_2_ (ml/hr) (n=7-10, one-way ANOVA, p<0.05) and total energy expenditure (kCal/hr) (n=7-10, one-way ANOVA, p<0.05) (Fig. 4K and 4L). *Posthoc* comparison showed energy expenditure to be significantly reduced in Cre-SEFL group in compared to other groups in the light cycle (n=7-10, Tukey’s multiple comparison test, p<0.05). *Posthoc* analysis confirmed that O_2_ consumption was reduced in the Cre-SEFL group compared to Cre-No SEFL group in the light phase (n=7-10, Tukey’s multiple comparison test, p<0.05). There were no changes in fat mass and lean mass, however, rectal temperature in Cre-SEFL were slightly higher than controls despite increase in EE. We also measured other parameters like food consumption and total locomotion (m/hr) and saw no main effect (n=7-10, one-way ANOVA, p>0.05) (Supplemental Fig. 4H and 4I). We also noticed decrease in VO_2_ and EE upon cold exposure (Supplemental Fig. 4J).

**Figure 4.**
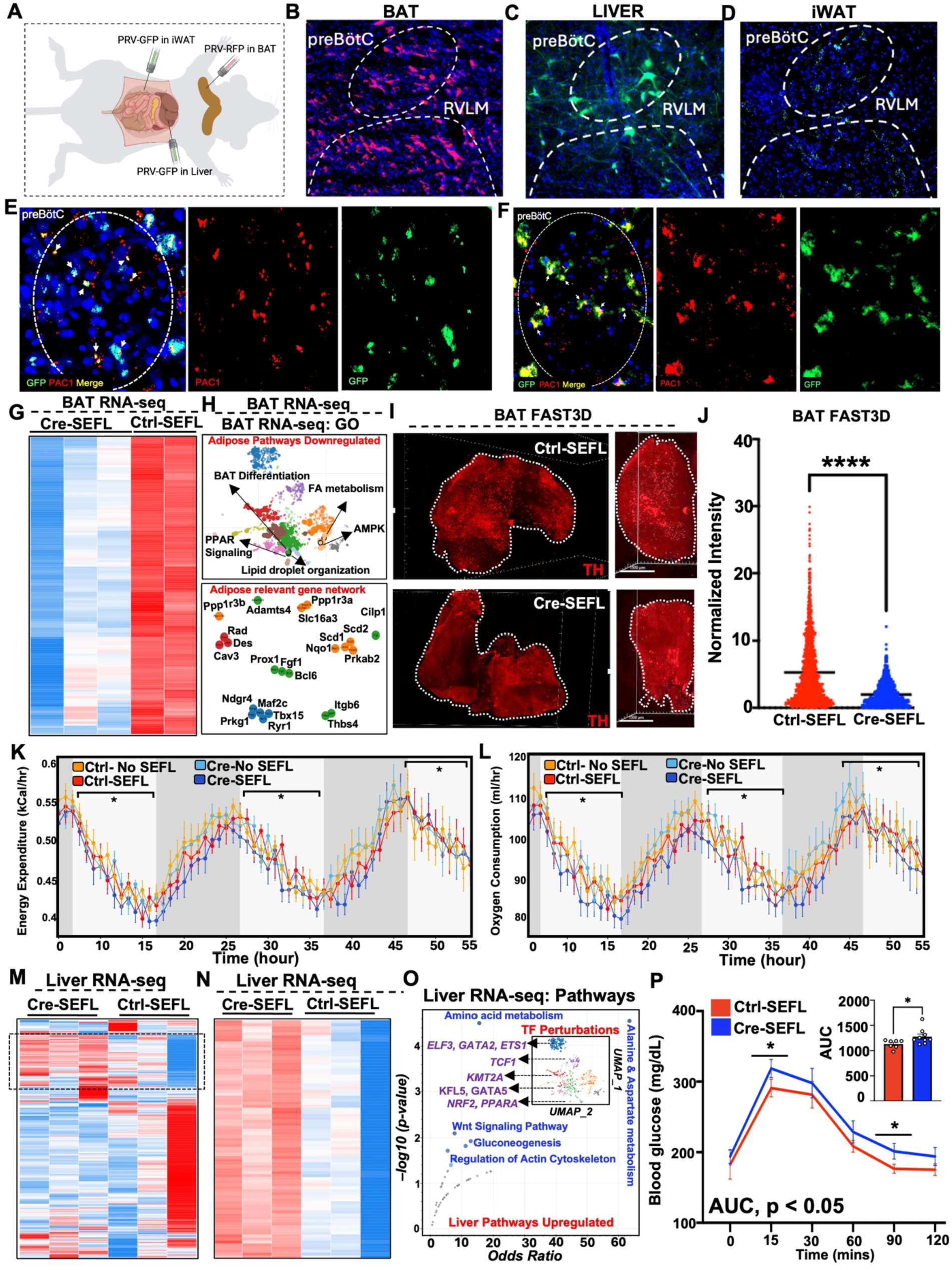
PAC1R deletion paired with SEFL disrupts glucose regulation and metabolic function. (A) A) Schematic of the retrograde tracing strategy. B) Representative images showing RFP expression in the preBötC following retrograde labeling of PRV-RFP in the BAT. C) Representative images showing GFP expression in the preBötC following retrograde labeling of PRV-GFP in the larger lobe of the liver. D) Representative images showing GFP expression in the preBötC following retrograde labeling of PRV-GFP in the iWAT. E and F) Representative RNAscope FISH multiplexed with IHC images showing PAC1r mRNA (red) and PRV-GFP (green) expression in the preBötC retrograde tracing from BAT (E) and liver (F). G) Heatmap showing downregulated genes in Cre-SEFL compared to Ctrl-SEFL from BAT RNA-seq analysis. H) Downregulated pathway and gene network analysis from Cre-SELF BAT RNA-seq data. I) Representative images of TH staining in the BAT from Ctrl-SEFL and Cre-SEFL mice. G) Quantification of TH signal intensity normalized to tissue volume in Ctrl-SEFL and Cre-SEFL group (n=2, unpaired t-test, t=19.92, p<0.0001). K and L) Energy expenditure and oxygen consumption metabolic chamber data from indicated mouse groups with 60 hours of measurement (n=7-10, one-way ANOVA, p<0.05). M and N) Heatmap of DEGs from liver RNA-seq with rectangle highlighting genes that are upregulated in Cre-SEFL. O) Pathway analysis and transcription factor gene network that are upregulated in Cre-SEFL compared to Ctrl-SEFL. P) Blood glucose levels during GTT in Ctrl-SEFL and Cre-SEFL mice at multiple time points following glucose injection (n=7-9, two-way ANOVA, F(1,84)=5.86, p=0.017); <u>Area under the curve (AUC) quantification of</u> glucose <u>response (n=7-9, unpaired t-test, t=2.235, p=0.0422)</u>.

To determine how loss of pre-Botz-PAC1 signaling controls liver biology under chronic stress, we performed bulk RNA-seq on whole-liver extracts collected 24 h after the SEFL session and visualized the data in heatmaps and a two-dimensional UMAP projection (Fig. 4M-4O).The left heatmap displays the top 150 genes assigned to down-regulated pathways (blue gradient). Two discrete modules emerged: (i) a broad suppression of oxidative-metabolism genes—including Cpt1a, Acadl, and Hadha—that paralleled the reduction in whole-body energy expenditure, and (ii) a coordinated decrease in lipid-handling transcripts (Apoa4, Fabp1, Mttp) consistent with reduced fatty-acid turnover. In sharp contrast, the middle heatmap highlights genes selectively up-regulated in Cre-SEFL livers (red gradient). These genes cluster into three functional axes: amino-acid catabolism (Tat, Got1, Oat), canonical gluconeogenesis (Pck1, G6pc), and Wnt/β-catenin signaling (Axin2, Ctnnb1, Tcf7l2), indicating a metabolic switch from lipid to amino-acid substrates and a potential proliferative/repair response. Dimensional reduction placed Cre-SEFL samples on a unique trajectory along UMAP_2, orthogonal to both control groups, underscoring the magnitude of PAC1-dependent remodeling. Pathway score overlays (grey dashed contours) confirmed that vectors pointing toward Cre-SEFL coordinates were enriched for amino-/aspartate-metabolism and liver-specific proliferative signatures, whereas vectors toward controls were enriched for oxidative and PPAR-regulated lipid pathways. Integrated differential TF analysis pinpointed a cluster of TFs whose motif occupancy changed most dramatically in Cre-SEFL livers: activators ELF3, GATA2, ETS1, TCF1, KMT2A and repressors KLF5, GATA5, NRF2, PPARA (black arrows). Notably, ELF- and GATA-family activation is often linked to hepatic inflammatory or regenerative programs, whereas loss of NRF2 and PPARA activity is compatible with diminished antioxidant and lipid-oxidation capacity, respectively. Together, the heatmaps and UMAP analysis reveal that, in the absence of preBötC-PAC1 signaling, chronic psychological stress precipitates a re-wiring of liver metabolism: lipid-oxidative circuits are perturbed, while amino-acid catabolism, gluconeogenic flux, actin-cytoskeletal remodeling, and Wnt signaling are up-regulated. TF-perturbation profiling implicates a shift from PPARA/NRF2-centric lipid control toward ELF/GATA-mediated inflammatory and proliferative pathways, highlighting a compensatory—yet potentially maladaptive—hepatic response to systemic metabolic stress in PAC1-deficient animals. Importantly, these changes were not attributed to the sympathetic tonality of livers, as we found no difference in TH staining in liver between Cre-SEFL and Ctrl-SEFL (Supplemental Fig. 4K-4M)

To investigate central mechanisms of stress-related metabolic adaptation in liver, we performed metabolic assays including, glucose tolerance tests (GTT), pyruvate tolerance tests (PTT), and insulin level assessments. Furthermore, we found that the Cre-SEFL group exhibited elevated blood glucose levels during the GTT compared to GFP-SEFL mice, with a significant main effect of PAC1R deletion (n=7-9, two-way ANOVA, F(1,84)=5.86, p=0.017) (Fig. 3E). *Post hoc* analysis revealed a significant increase in blood glucose specifically at 15 and 90 minutes after glucose injection (n=7-9, Tukey’s multiple comparison test, p<0.05). Pyruvate serves as a key substrate for gluconeogenesis; therefore, to assess the liver’s ability to convert pyruvate into glucose, we performed the PTT. Analysis revealed no significant effect of PAC1R deletion on blood glucose levels following pyruvate injection (n=7-9, two-way ANOVA, F(1,78)=0.205, p=0.65) (Fig. 3A). Additionally, no significant differences in plasma insulin levels were observed between the Ctrl-SEFL and Cre-SEFL groups (n=7-9, two-way ANOVA, F(1,35)=1.449, p=0.24; AUC: n=7-9, unpaired t-test, t=0.117, p=0.908) (Supplemental Fig. 3B). We also performed an insulin tolerance test (ITT) and observed no group effects on insulin sensitivity (n=5, two-way ANOVA, F(1,42)=0.019, p=0.89) (Supplemental Fig. 3C).

## Discussion

Our work, along with that of many others in the field of stress (de Sousa Abreu et al., 2024; Garfinkel and Critchley, 2016; Garfinkel et al., 2014; Hsueh et al., 2023; Klein et al., 2021; Yoshimoto et al., 2024), has contributed to a growing shift in focus —moving beyond a traditionally brain-centric framework to encompass a more integrative perspective that includes key physiological mechanisms like respiration. This underscores the importance of mechanistically dissecting how respiratory function contributes to the regulation of stress behaviors and the emergence of related metabolic dysfunction. In our study, we demonstrated that disrupting breathing rhythms through ablation of PAC1R in the preBötC leads to heightened fear sensitization in animals previously exposed to stressors, regardless of the intensity of the stressor. We further established a tight coupling of cardio-respiratory responses as shown by increased ⊗HR during bouts of freezing in animals and anesthetized BRPM with ablated PAC1R. Anatomical tracing revealed neuronal projections from the preBötC to both BAT and the liver, suggesting a novel respiratory-metabolic regulatory pathway. Functionally, PAC1R ablation in the preBötC combined with SEFL led to glucose intolerance, disrupted energy expenditure, stress-induced hyperthermia, and reduced sympathetic innervation of BAT.

Our SEFL paradigm reliably captures both associative and non-associative components of fear learning (Nishimura et al., 2022; Perusini et al., 2016; Poulos et al., 2015; Rau et al., 2005) , and effectively models key features of PTSD, including avoidance, hyperarousal, hypervigilance, and an exaggerated startle reflex—core indicators of autonomic dysregulation. By selectively ablating PAC1R in the preBötC, we disrupt normal breathing patterns and induce a state of autonomic imbalance, characterized by increased ⊗HR and BRPM in Cre groups. This disruption appears to drive both non-associative fear generalization and fear sensitization. Behaviorally, this is evidenced by increased fear generalization to a novel context (Context B) in Cre-SEFL mice, as well as elevated freezing responses on Day 3 in Cre-No SEFL animals following a single exposure to footshock in Context B. Furthermore, to address the ceiling effect observed with higher-intensity stress, we reduced the Day 1 stressor to five 1 mA footshocks. Even under these conditions, the Cre-SEFL group exhibited significantly increased freezing on Day 3 compared to the Ctrl-SEFL group, highlighting the heightened sensitivity induced by PAC1R ablation in the preBötC. These findings strengthen our hypothesis that elevated respiratory rate contributes to the amplification of maladaptive fear learning. However, our current studies do not yet capture the precise temporal or mechanistic correspondence between respiratory dynamics and behavioral output. The lack of high-resolution, real-time measurements linking specific breathing patterns to discrete fear-related behaviors limits our ability to determine causality.

Our spatial transcriptomic analysis of the preBötC following PAC1R deletion revealed a coordinated reorganization of transcriptional programs, with downregulation of genes essential for cholinergic neurotransmission (Ache, Slc5a7, Gnas, Syp, Ptprn), inhibitory tone (Dlk1, Kcnk9, Gch1, Syt4), and synaptic vesicle cycling, alongside upregulation of excitatory and metabolic regulators in glutamatergic neurons (Oxct1, Pde4b, Trpc3, Itgb8). These changes point to a shift in excitatory–inhibitory balance and neuromodulatory capacity within the preBötC, a respiratory hub that also interfaces with autonomic premotor circuits. The systemic phenotype of PAC1R deletion marked by reduced brown fat innervation, diminished energy expenditure, and altered hepatic metabolism, suggests that these central molecular alterations may translate into disrupted downstream autonomic output.

The novelty of our study lies in the discovery of significant metabolic disruption resulting from PAC1R ablation in the preBötC. Through GTT (Andrikopoulos et al., 2008; Hahn et al., 2024; Virtue and Vidal-Puig, 2021), we observed a clear impairment in glucose regulation in animals with targeted PAC1R ablation combined with SEFL. Interestingly, despite the pronounced glucose intolerance, these animals showed no significant changes in circulating serum insulin levels or insulin sensitivity (Hahn et al., 2024; Virtue and Vidal-Puig, 2021), suggesting that insulin production and peripheral insulin responsiveness remain largely intact. Furthermore, the preservation of normal pyruvate tolerance (Hahn et al., 2024) responses indicates that increased hepatic gluconeogenesis is unlikely to be the underlying cause of the observed glucose dysregulation. We also examined whether increased fat mass (Williams et al., 2014) or elevated corticosterone levels (Andrews et al., 2002; Fichna and Fichna, 2017; Kuo et al., 2015) could account for the observed glucose intolerance, but found no significant differences in either parameter. Taken together, these findings suggest that PAC1R ablation in the preBötC could potentially lead to disrupted beta-cell function or impaired glucose uptake at a cellular level, rather than systemic insulin, hepatic, or corticosterone-related dysfunction. These findings highlight the need for further studies to elucidate the specific mechanisms underlying glucose intolerance in PAC1R-deficient mice. Moreover, we observed increased rectal temperature immediately following SEFL and low energy expenditure at later time points, which together suggest abnormalities in thermoregulation (Abreu-Vieira et al., 2015) and potential stress-induced hyperthermia (Blenkuš et al., 2022; Van der Heyden et al., 1997), respectively. While the acute rise in temperature is likely driven by stress, the long-term effect of PAC1R ablation combined with SEFL appears to result in a sustained decrease in energy expenditure during the light phase. This effect is likely due to reduced sympathetic innervation of BAT (Bamshad et al., 1999; Bartness et al., 2010), as supported by our identification of a novel neuronal projection from PAC1-expressing neurons to BAT. These findings point to a central, PAC1R-dependent mechanism that regulates thermogenesis and energy balance via modulation of sympathetic output to BAT.

## Methods

### Animals

All the experimental procedures were conducted following the guidelines set by the Institutional Animal Care and Use Committee (IACUC) at the Icahn School of Medicine at Mount Sinai. A total of approximately 60 ten-week-old PAC1R floxed mice, and 10 ten-week-old ADCYAP-GFP mice, both males and females, were obtained from Jackson Laboratories. Mice were housed in a temperature-controlled room under a 12-h light–12-h dark cycle and under pathogen-free conditions. Mice were fed a chow diet (4.5% fat/calorie). At the time of euthanization, tissues were collected after decapitation, frozen immediately, and stored at -80°C.

### Virus

Mice were either injected with AAV2-hsyn-GFP-Cre for Cre-mediated deletion of PAC1R or AAV2-hsyn-GFP for control animals (The UNC vector core). To identify brain regions sending monosynaptic PACAPergic inputs to the preBötC, we injected Cholera toxin B conjugated with Alexa 647 (CTB-A647) (Thermo Fisher Scientific) in the preBötC of ADCYAP-GFP mice. To identify brain sites sending the projections to liver, BAT and iWAT, we used a transsynaptic tracing technique with a pseudorabies virus strain, PRV-GFP (PRV-152; NIH CNNV- University of Pittsburgh).

### Cre-mediated PAC1R deletion

To ablate PAC1R receptors in neurons, mice were secured in a stereotaxic apparatus under 2% isoflurane anesthesia and injected with rAAV2-hsyn-GFP-Cre to achieve neuronal deletion of PAC1R receptors. Control mice were injected with rAAV2-hsyn-eGFP. Mice were injected with 0.5 μl of virus bilaterally into the preBötC using coordinates (LM: ±1.25; AP: −6.78; DV: -5.70 from Bregma). Virus was microinfused into the preBötC with a 10 μl Hamilton Syringe fitted with a 1 mm glass pipette with no filament at a rate of 0.2 μl/min. After completion of infusion, the glass pipette was left in position for 10 min to allow for diffusion. Using the glass pipettes that are commonly used for electrophysiological recordings with single-cell resolution allowed for confining the virus infusion to the preBötC. Using this method, we have previously targeted structures that are as small as the preBötC (Rajbhandari et al., 2021). Immediately after surgery, mice were given ad libitum access to a 0.5 mg/kg Sulfamethoxazole solution in drinking water for 5 days. The antibiotic regimen is standard procedure to prevent infections after surgeries. Since this is administered for 5 days after surgeries and the experimental procedure do not start until 3 weeks later, we do not expect it to have effects on metabolic functions. Mice also received a subcutaneous injection of the anti-inflammatory drug Rimadyl (5 mg/kg) immediately after, and 1 day following surgery. We allowed for 21 days of viral transfection after surgery before behavioral testing.

### CTB injections

To trace the PACAPergic inputs to the preBötC, we used a monosynaptic tracing strategy using CTB-A647. Mice were injected with 0.5 μl of virus bilaterally into the preBötC using coordinates (LM: ±1.25; AP: −6.78; DV: -5.70 from Bregma). Virus was microinfused into the preBötC with a 10 μl Hamilton Syringe fitted with a 1 mm glass pipette with no filament at a rate of 0.2 μl/min. After completion of infusion, the glass pipette was left in position for 10 min to allow for diffusion. Mice also received a subcutaneous injection of the anti-inflammatory drug Rimadyl (5 mg/kg) immediately after, and 1 day following surgery. The mice were also given ad libitum access to a 0.5 mg/kg Sulfamethoxazole solution in drinking water for 5 days. Mice were euthanized 10 days after the stereotaxic injection.

### PRV injections

All virus injections were performed according to Biosafety Level 2 standards. Mice (n=4, 2 for liver, 2 for BAT) were anesthetized with 0.5mg/kg of ketamine/xylene. The PRV was injected into the left lobe of the liver and BAT (10 × 100 nL). After injection, the needle was left in place for 15 seconds. The skin was sutured closed, and analgesia (Rimadyl 5 mg/kg) was administered via i.p. injections for the first two days.

### Stress-enhanced fear learning

SEFL is a robust and powerful rodent model that recapitulates many clinical aspects of PTSD. This model was established over a period of 3 days (Nishimura et al., 2022; Rajbhandari et al., 2018). On day 1, fear conditioning chambers were set up to serve as Context A in which the mice received the footshock stressors. In this context, the mice were directly transported to the chambers from their home cages. The chambers contained A-frame inserts and a level grid floor. Within the chamber, the conditional stimulus was presented in the form of white light, white noise, and the olfactory cue of a wet paper towel. All the mice were placed in the chamber for 1 hour, and 10 random foot shocks of 1 mA were administered during this period. On day 2, we subjected all mice to a mild stressor (1-foot shock) in context B and assessed their fear response to this stressor. For this context, mice were individually transported in opaque containers to the behavior room. To modify the chamber environment, an additional light source was introduced, a staggered grid floor and curved wall insert were added, and 1% acetic acid served as the olfactory cue. On day 2, the mice were placed in the chambers for a period of 4 minutes and 30 seconds. The mice received the 1-foot shock of 1 mA at the end of the 4th minute to serve as a mild stressor. On day 3, the mice were placed back in context B for 8 minutes, and the fear response to the mild stressor in context B was assessed by measuring freezing behavior. The percentage of freezing, defined as lack of movement, was calculated before every footshock on day 1, for the first 4 minutes on day 2, and 8 minutes on day 3. An additional measure of shock reactivity was calculated during the footshocks on day 1, and the 4th-minute shock on day 2 using the same VideoFreeze system.

### Measure of Freezing and Shock Reactivity

Freezing is a complete lack of movement except when the animal is breathing. Freezing was measured using VideoFreeze™ Video Fear Conditioning Software by Med Associates, designed to capture and freeze video frames. The software measures freezing by measuring the total time spent motionless during the session. The threshold for activity detection was set at 1 au. The animal was considered to be freezing when the activity score remained below this threshold for a minimum freeze duration of 10 frames (0.33 seconds). Percentage freezing = freezing time/total time×100 for the period of interest. The data are presented as the mean percentages (+/− SEM). Shock reactivity is an index of pain reactivity to the incoming foot shock. The activity burst velocity is a reliable predictor of foot-shock intensity and may be a useful indicator of pain reactivity. We calculate shock reactivity by the motion index during the 1-s period during the shock on Day 2. The motion index in cm/s (+/− SEM) was computed for the period of interest.

### Open Field Light Gradient Anxiety Test

The mice were also tested for anxiety-like behavior in a modified open-field task that incorporated a light gradient (Nagai et al., 2019). The open field light gradient test was used to measure anxiety-like phenotypes using parameters such as distance traveled, and time spent in the dark zone. A rectangular, white, translucent polyethylene box (46 × 86 × 30 cm) was placed in a dark room. Three lamps were placed on one end of the box. A camera suspended above captured the animal’s activity. The test was divided into three phases during the 12-minute run. During the first four minutes, the lights are turned off, and the animal can explore the open field in the dark phase 1. In the next four minutes, the lamps are turned on, and a light gradient is created within the open field. The zone closest to the light source, called the “light zone”, has an illumination index of ∼120 lux. The “middle zone” has intermediate illumination of ∼50 lux, and the zone farthest from the light source, the “dark zone”, has an illumination index of ∼10 lux. In the last phase of the experiment, dark phase 2, the lights are turned off again for four minutes. The locomotive activity and time spent in the dark zone were calculated using EthoVision XT software.

### Elevated Plus Maze

The EPM is a behavioral task commonly used to screen for anxiety-like behavior. The maze is gray in color, and consists of two perpendicular intersecting runways (5 cm wide X 35 cm long) connected by a central area (5 × 5 cm). One runway has tall walls (closed arms; 15 cm in height) while the other runway has no walls (open arms). The maze was elevated 40 cm from the floor. Mice were placed in the center of the maze and were allowed to explore the maze for a total of 5 min under controlled lighting conditions (∼35 lx) in a behavioral testing room. The cumulative frequency of entries in the open arms (s), and time spent in the center (s) was calculated using EthoVision XT software.

### Telemetric heart rate measurement

Heart rate was recorded with the MouseOx Pulse oximeter (s-collar clip, shaved neck; Starr Life Sciences) following the manufacturer’s instructions. Mice were shaved around the neck and acclimated to moving with the collar sensor for at least six days during the standard handling procedure. Heart rate was recorded during Day 2 and Day 3 of the SEFL protocol. Heart rate deviations (ΔHR) from baseline were calculated by first binning raw heart rate data into 1s bins, and then building a z-score of the whole trace using the mean and SD of the first 100s of each experimental session considered as baseline (Klein et al., 2021).

### Anesthetized telemetric breath rate measurement

Breathing was recorded using the MouseOX Pulse oximeter. Post SEFL, shaved mice were placed under 1% isoflurane and breath rate per minute (BRPM) was measured for 5 minutes.

### Internal body temperature measurement

Internal body temperature was measured using rectal temperature probe at baseline; one day prior to SEFL, during SEFL; immediately after SEFL day 3, and post-SEFL; a week after SEFL day 3 for 4 hours, with measurements taken every half hour. The temperature of each mouse was measured sequentially by inserting a thermistor probe 2 cm into the rectum. The probe was dipped into petroleum jelly before inserting into the rectal cavity and was held into the rectum till a stable rectal temperature was measured for 10 s.

### Indirect calorimetry using a metabolic chamber

The mice were placed in an indirect calorimetry chamber for 72 hours to study metabolic function after 1 week of SEFL. The data collected via indirect calorimetry included oxygen consumption, carbon dioxide production, energy expenditure (EE), the respiratory exchange ratio (RER), energy balance, food and water intake, locomotor activity, and body mass. Body weight was considered a covariate in the statistical analysis for some indirect calorimetry measurements in the metabolic chambers, such as oxygen consumption, carbon dioxide production, EE, and food and water intake. The RER and locomotion were analyzed without a covariate, as they are known to be independent of body weight. This information highlights changes in metabolic physiology in PAC1R No-deletion/ PAC1R deleted animals after receiving either No-SEFL or SEFL. We analyzed the data with CalR (Mina et al., 2018), which considers activity, food intake, and other parameters, allowing us to derive accurate indirect calorimetry values.

### Glucose tolerance test

For the glucose tolerance test, two days after the mice were taken out of the metabolic chambers, the mice were fasted for 4 hours. The baseline blood glucose levels (mg/dL) of these mice were measured using one drop of blood collected from the tip of the tail and a glucometer. For the test, 1 g of glucose/kg body weight was injected intraperitoneally (i.p.) into each mouse. Blood glucose levels were determined at 0, 15, 30, 60, 90, and 120 min.

### Pyruvate tolerance test

Mice were fasted for 6 h prior to the test. The baseline blood glucose levels (mg/dL) of these mice were measured using one drop of blood collected from the tip of the tail and a glucometer. Sodium pyruvate (2.5 g/kg body weight) was injected intraperitoneally (i.p.) into each mouse. Plasma glucose was measured in blood samples collected from the tail at 15, 30, 60, 90, and 120 min after injection of sodium pyruvate by a glucometer.

### Insulin tolerance test

Mice were fasted for 6 h prior to the test. Baseline blood glucose levels (mg/dL) of mice were measured using one drop of blood collected from the tip of the tail using a glucometer. Human insulin (1U/kg body weight) was injected intraperitoneally (i.p.) into each mouse. Plasma glucose levels were measured from the tail blood at 15, 20, 60, 90 and 120 min after injection of insulin.

### Serum sample collection for ELISA

Mice were fasted for 4 hours before blood collection from the tail vein. Blood was collected into an EDTA-coated tube, centrifuged at 5,000 rpm for 10 minutes at 4°C, and the plasma was separated and stored at −80°C until further analysis. Plasma insulin levels were measured at baseline, 10 minutes, and 30 minutes after the intraperitoneal injection of 1 g glucose/kg body weight using ELISA (Mercodia #10–1247-01). Corticosterone levels were measured at baseline using ELISA (Crystal Chem #80556), following the manufacturer’s protocol.

### Immunohistochemistry

For immunohistoschemistry labeling, 40 μm coronal sections were cut from mice brains (n=5). Sections containing the preBötC were blocked and permeabilized in a solution containing PBS + 3% normal goat serum + 0.1% Triton X-100 for 1 h. An anti-cFos primary antibody was diluted 1:2000 in PBS + 3% normal goat serum + 0.1% Triton X-100, and sections were incubated overnight at 4°C, then washed in PBS, and subsequently incubated with an anti-rabbit AlexaFluor-488 secondary antibody diluted 1:500 for 3 hours at room temperature. Sections were washed in PBS and cover-slipped using Prolong Gold mounting media (Thermo Fisher Scientific) with 4′,6-diamidino-2-phenylindole (DAPI), and the edges were sealed with clear nail polish. For the retrograde tracing from the preBötC using CTB-A647, 40 μm coronal brain sections containing the brain regions known to project to the preBötC was collected, and endogenous fluorescence was detected by cover-slipping with Prolong Gold (Thermo Fisher Scientific) with DAPI, and the edges were sealed with clear nail polish. For cFos quantification and detection of retrograde CTB-A647 expression, sections were imaged using a 20X objective on a Zeiss AxioImager Z2M with ApoTome. cFos expression was analyzed using FIJI software by counting the number of cFos labeled with GFP within a standardized region of the preBötC.

### RNAscope in situ hybridization

Brain tissues were sectioned at 20 μm using a cryostat at −20°C, and slices containing the preBötC were collected on Superfrost Plus microscope slides (Thermo Fisher Scientific) and stored at −80°C. PAC1R receptor deletion was confirmed using RNAscope to analyze RNA expression in tissue sections (ACD Biotechne). Briefly, in situ hybridization was performed following the RNAscope® 2.5 HD Assay - RED protocol for fresh frozen sections. After labeling, sections were cover-slipped using Prolong Gold (Thermo Fisher Scientific) with DAPI, and the edges were sealed with clear nail polish. For PAC1R mRNA quantification, sections containing the preBötC were imaged using a 20X objective on a Zeiss AxioImager Z2M with ApoTome. PAC1R mRNA expression was analyzed using FIJI software by measuring the mean fluorescence intensity of PAC1R mRNA labeled with mCherry within a standardized region of the preBötC.

### RNAscope in situ hybridization multiplexed with Immunohistochemistry

To visualize the colocalization of PAC1R receptors on GABA(GAD2) or glutamatergic (Vglut2) neurons, and for the visualization of PAC1R receptor colocalization with the RFP expression from retrograde PRV-RFP injections in the periphery, we followed the dual ISH and IHC strategy. We followed the RNAscope ISH protocol as detailed in the section above, following which IHC staining was carried out to detect protein expression, using antibodies specific to the protein of interest as described in the section “Immunohistochemistry” using the ACD Bio Multiplec reagent kit.

### Fast 3D tissue clearing

Fast 3D tissue clearing was performed as previously described (Kosmidis et al., 2021). Briefly, tissue samples were dehydrated and delipidated using tetrahydrofuran (THF). THF treatment induces a linear expansion of the tissue relative to its original size, after which the samples were washed with distilled water. To enhance the endogenous signal, the original protocol was modified to include immunostaining of cFos or the virally conjugated fluorescent markers. Finally, optical clearing was performed using the LifeCanvas EasyIndex solution, designed to match the refractive index and improve tissue transparency.

### Microscopy

The tissue sections were analyzed using a Zeiss AxioImager Z2M with ApoTome microscope. Images were analyzed with Fiji image processing software (NIH, Bethesda, MD, USA; RRID:SCR_002285). Mean fluorescent intensity of PAC1R RNA was measured on sections of preBötC. The cleared brain, liver, and BAT samples were imaged using the Lifecanvas SmartSPIM lightsheet microscope with a 1.6x objective for liver samples with dynamic focus and a step size of 4 μm, and a 3.6x objective was used for BAT and brain samples imaging with dynamic focus and a step size of 2 μm. Images were analyzed using Imaris (Oxford Instruments; RRID:SCR_007370).

### Bulk RNA-sequencing

Total RNA was prepared as described (Tong et al., 2016). Strand-specific libraries were generated from 500 ng total RNA using the TruSeq Stranded Total RNA Library Prep Kit (Illumina). cDNA libraries were single-end sequenced (50bp) on an Illumina HiSeq 2000 or 4000. Reads were aligned to the mouse genome (NCBI37/mm9) with TopHat v1.3.3 and allowed one alignment with up to two mismatches per read. mRNA RPKM values were calculated using Seqmonk’s mRNA quantitation pipeline. All FPKMs represent an average from three biological replicates for in-vitro studies for library construction. A gene was included in the analysis if it met all of the following criteria: The maximum FPKM reached 10 at any time point, the gene length was > 200bp as determined by the DESeq2 package in R Bioconductor. P values were adjusted using the Benjamini-Hochberg procedure of multiple hypothesis testing (Benjamini and Hochberg, 1995).

### Sample preparation for 10x Visium HD Spatial Assay

Mice were anesthetized with 1% isoflurane, after which their brains were extracted and flash-frozen in isopentane cooled with dry ice. The brains were quickly dissected and embedded in OCT compound. Proper orientation was ensured before freezing the samples in a dry ice/ethanol bath. Brains were then stored at −80 °C until cryosectioning.

### 10x Visium HD Spatial Data Analysis

10X Genomics’ Space Ranger software was used to pre-process the sequencing data and align them to the mm10 reference transcriptome. The output of Space Ranger was the input for the subsequent steps. All samples underwent identical quality control and processing steps. Downstream spatial transcriptomics analysis including normalization, integration, and clustering was performed by Seurat (v.5) using Feature barcoded expression matrices as input (Hao et al., 2024). The well-performing popular single-cell transcriptomics analysis, the Seurat pipeline has been extended to reliably handle spatial transcriptomics data. Seurat enables excellent data normalization, dimensionality reduction, clustering and strong visualization. The Seurat (v.5) was used with the default parameters as described in the spatial clustering vignette. In brief, expression matrices were read into the R environment, and a Seurat object was created for each sample. “FindClusters” function was ran at different resolutions and the best resolution giving the most desired number of clusters was chosen. A few options were employed for visualization of the clusters. (1) “DimPlot” function to generate the traditional UMAP plot showing the clustering results. (2) “SpatialDimPlot” to show the distribution of identified clusters by overlaying them onto the tissue image and to highlight the location of selected clusters. Basically, Seurat clusters cells use a graph-based method that begins with a K-Nearest Neighbors graph based on the PCA space. The cells are then clustered using the Louvain algorithm, and finally, a non-linear dimensional reduction is carried out using UMAP. To visualize all spots in a two-dimensional plot, a UMAP was created with Seurat’s “RunUMAP” function using the first 30 principal components. Enriched genes of clusters were identified with Wilcoxon tests as implemented in the “FindMarkers” function. Well-known cell-type markers were used for assigning cell type labels to each cluster i.e. cell type annotations. According to Seurat, data from control and KO mice were annotated and combined into a merged object for downstream extensive analysis using “integrated.cca” for dimensional reduction. The combined object enabled us to further compare the cell type-specific changes, including the cell proportion change and differential gene expression analysis between control and KO mice. Volcano plots were also used to visualize differentially expressed genes among samples and specified clusters. Considered as the prebotc region, clusters 32 and 42 from KO mice (N) and 38 and 40 from control mice (R) were the primary cell group targeted for more in-depth analysis to elucidate their properties.

### Illustrations

Illustrations were generated with BioRender (www.biorender.com) under an academic license.

### Statistics and power calculations

The data are shown as the mean ± S.E.M. Significance was determined by a two-tailed unpaired t-test (parametric distribution), 2-way ANOVA with the Bonferroni post hoc correction, ANCOVA, Tukey’s multiple comparison test, or the Mann‒Whitney test (nonparametric distribution). The significance level was set at an alpha level of 0.05.

## Acknowledgements

P.R is supported by R01DK136035, DP1DK140003, Irma T. Hirschl/Monique Weill-Caulier Foundation Scholar Award. A.K.R. is supported by R21DK129908. The funders had no role in study design, data collection and interpretation, or the decision to submit the work for publication. We thank the Mount Sinai Genomics Core Facility for their assistance with spatial transcriptomics.

## Author’s contributions

S.S. conducted the behavioral experiments and metabolic studies; A.F. analyzed the spatial transcriptomics data; R.Y. conducted the RNAseq experiment; P.T. and S.J.D. carried out the behavior experiments and tissue harvest; D.E. helped with the peripheral retrograde injections; Y.J. and A.W. conducted the metabolic chamber experiments. S.S, P.R., and A.K.R. supervised the studies; P.R., and A.K.R. conceived and designed the studies and wrote the manuscript.

## Supplemental Figure Legends

**Figure 1.**
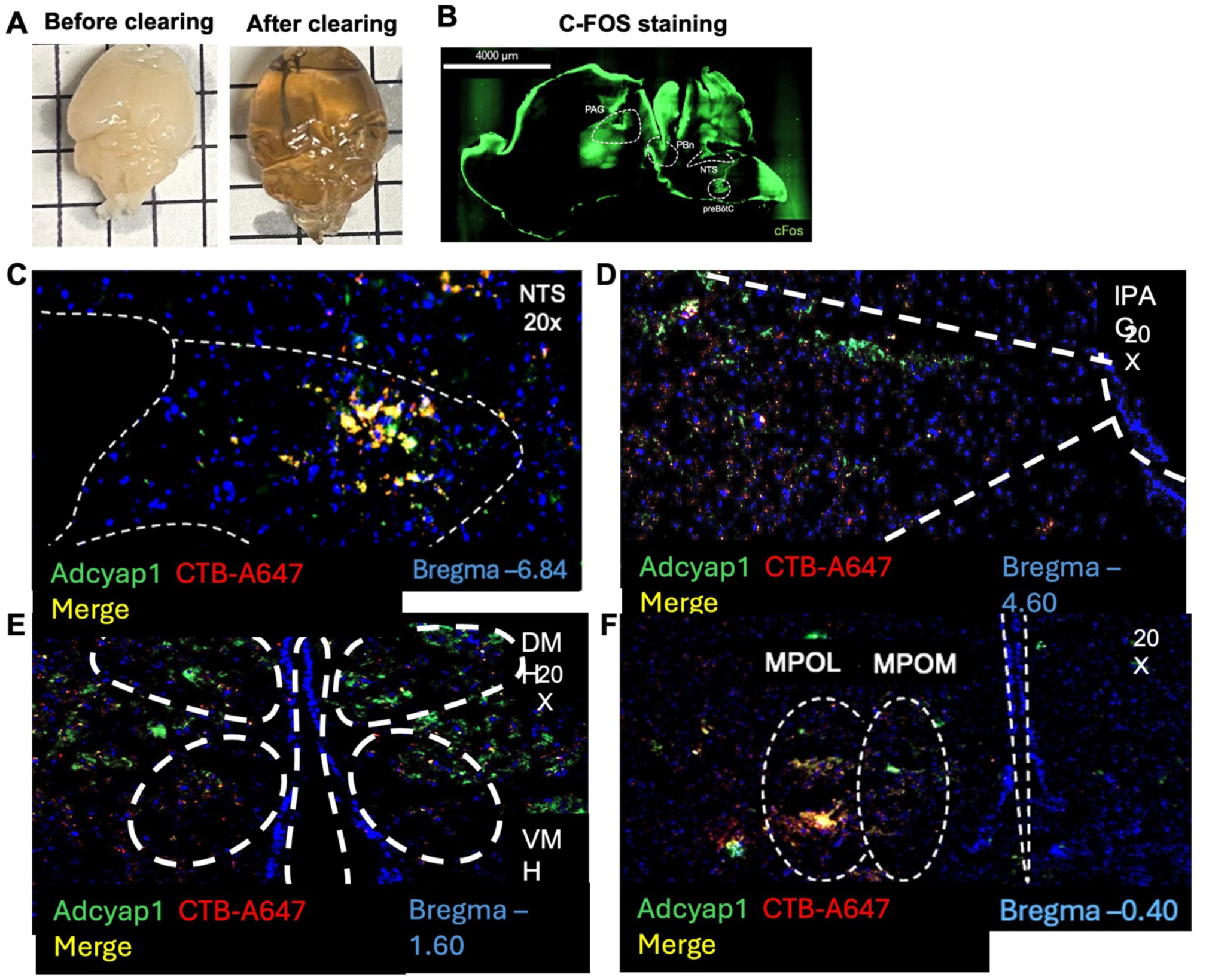
Whole brain cFOS staining and PACAPergic inputs to the preB<u>ö</u>tC. A) Image of brain harvested from SEFL-exposed mice before and after clearing. B) Representative image of whole-brain cFos immunostaining illustrating widespread neuronal activation following SEFL. C–F) Representative images of CTB-A647 labeling (red) in brain regions sending monosynaptic projections to the preBötC, showing colocalization with PACAP (green) in the medial nucleus of the solitary tract (mNTS; C), lateral periaqueductal gray (lPAG; D), dorsomedial hypothalamus (DMH; E), and medial preoptic area (MPO; F).

**Figure 2.**
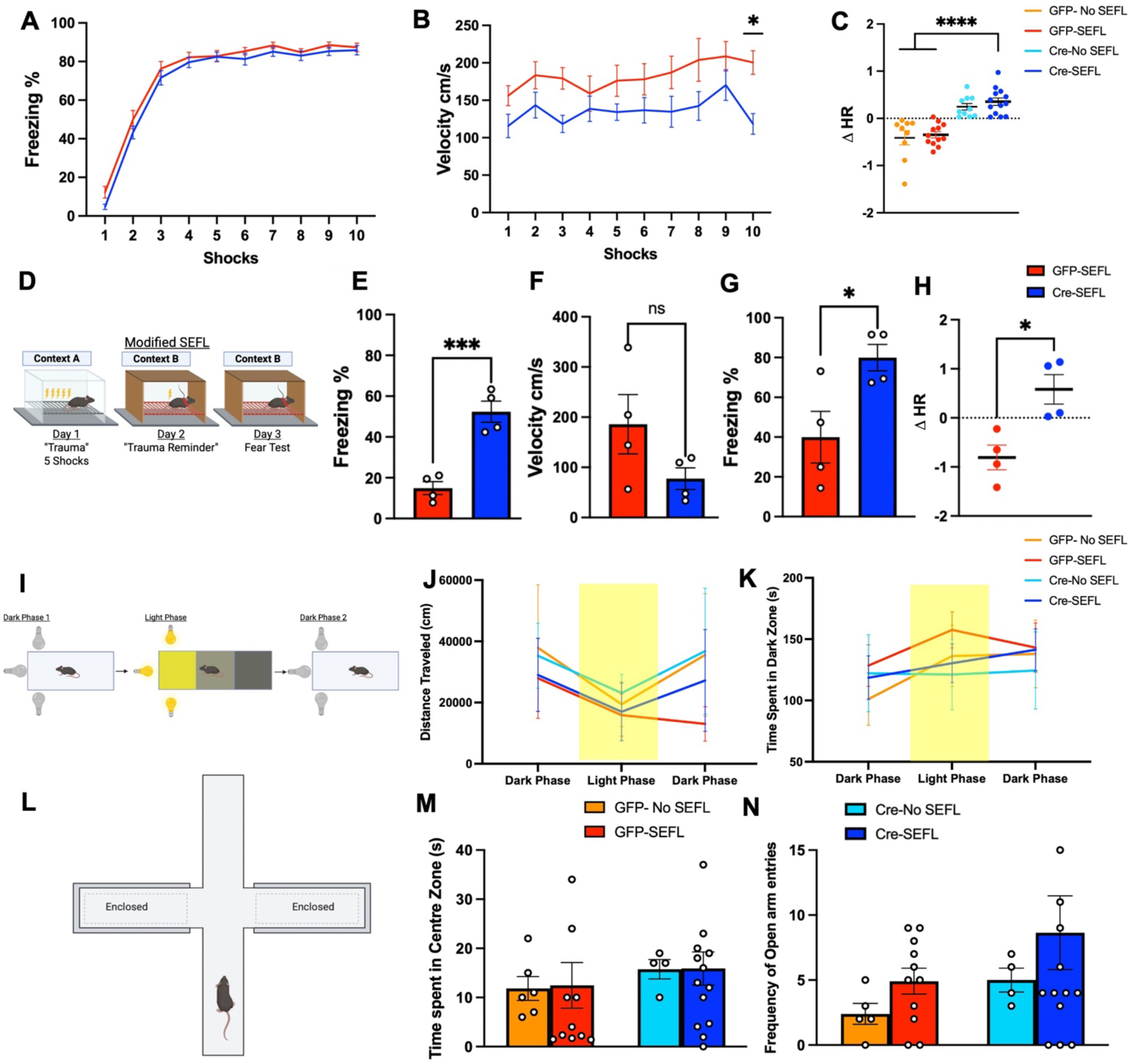
Ablation of PAC1R in the preBötC disrupts fear-related behavior without altering anxiety-like responses. A) Percent freezing during each successive footshock presentation on Day 1 of SEFL in Ctrl-SEFL and Cre-SEFL groups (n=18-20, two-way ANOVA, F(9,360)=0.3511, p=0.9659). B) Shock reactivity across successive footshock presentations on Day 1 of SEFL in Ctrl-SEFL and Cre-SEFL groups (n=18-20, two-way ANOVA, F(1,360)=34.17, p<0.0001). C) Average ΔHR during the entire duration of freezing on Day 3 of the SEFL paradigm in all four groups (n=9-13, one-way ANOVA, F(3,40)=19.13, p<0.0001). D) Schematic of modified SEFL protocol with reduced stressor footshocks on Day 1 of SEFL. E) Percent freezing measured on Day 2 of modified SEFL in Ctrl-SEFL and Cre-SEFL groups (n=4, unpaired t-test, t=6.135, p=0.0009). F) Shock reactivity measured via velocity (cm/s) in response to Day 2 footshock in the modified SEFL paradigm (n=4, unpaired t-test, 1.72, p=0.1362). G) Percent freezing on Day 3 of modified SEFL (n=4, unpaired t-test, t=2.733, p=0.0341). H) Average ΔHR of Ctrl-SEFL and Cre-SEFL group during bouts of freeezing on day 3 of modified SEFL (n=4, unpaired t-test, t=3.561, p=0.0119). I) Schematic of the open field light gradient task. J) Measure of distance traveled in cm over the three phases of the task: dark phase 1, light phase ,and dark phase 2 (n=10-19, two-way ANOVA, F(3,153) =0.5697, p=0.6358). K) Measure of the time spent in the dark zone of the rectangular open field during the three phases of the task (n=10-19, two-way ANOVA, F(3,153)=0.5564, p=0.6446). L) Schematic of the elevated plus maze task. M)Measure of time spent in the center zone of the elevated plus maze during the 5-minute trial (n=4-14, one-way ANOVA, F(3.31)=0.2573, p=0.855). N) Measure of time spent exploring the open arms of the elevated plus maze (n=4-14, one-way ANOVA, F(3,30)=1.125, p=0.3548).

**Figure 3.**
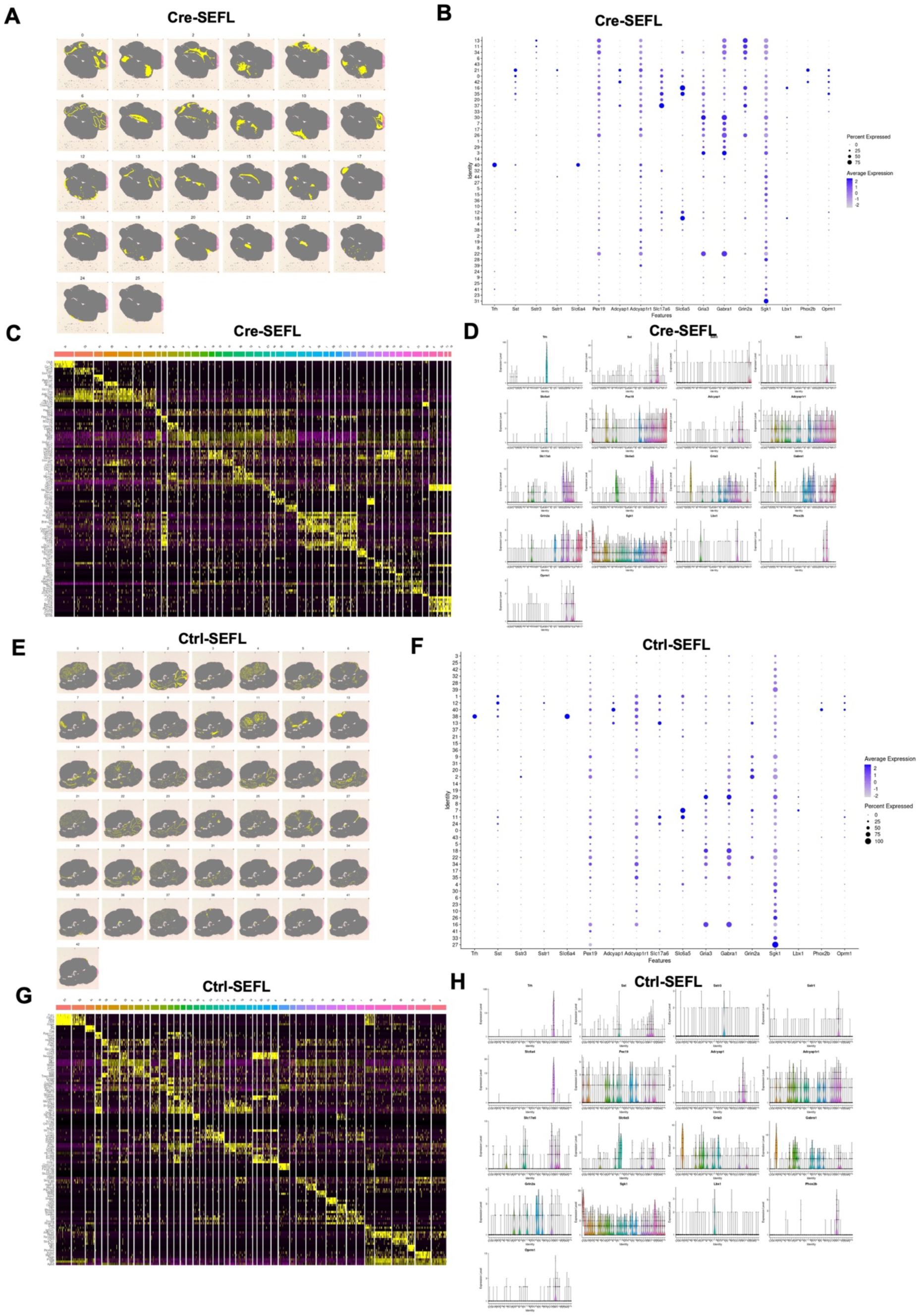
Spatial transcriptomics of brain stem slice. A) Spatial dimplot of brain regions from gene expression analysis from Cre-SEFL mice. B) Dot plot of genes expressed in annotated clusters from Cre-SEFL mice. C) Heatmap of genes expressed in each cluster from Cre-SEFL mice. D) Volcano plot of genes expressed in preBötC from Cre-SEFL mice. E) Spatial dimplot of brain regions from gene expression analysis from Cre-SEFL mice. F) Dot plot of genes expressed in annotated clusters from Cre-SEFL mice. G) Heatmap of genes expressed in each cluster from Cre-SEFL mice. H) Volcano plot of genes expressed in preBötC from Cre-SEFL mice.

**Figure 4.**
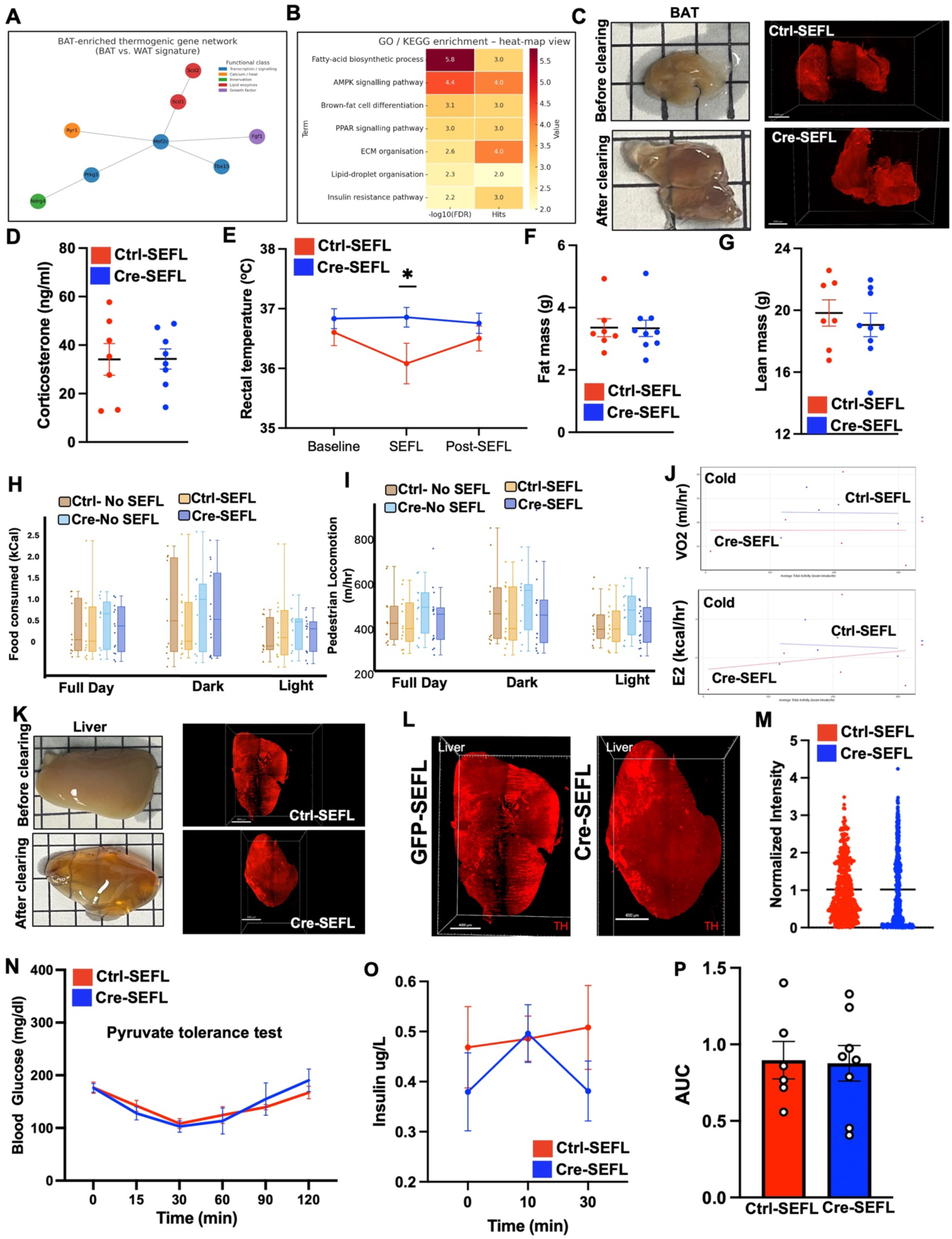
Metabolic and physiological consequences of PAC1R ablation in the preBötC. A) BAT-enriched thermogenic gene network analysis (BAT vs. WAT signature). B) GO/KEGG pathway enrichment heatmap showing significantly enriched metabolic pathways. C) Representative images of BAT before and after tissue clearing, with corresponding 3D reconstructions of TH immunolabeling in Ctrl-SEFL and Cre-SEFL groups. D) Plasma corticosterone levels in blood of Ctrl-SEFL and Cre-SEFL groups (n=7-8, unpaired t-test, t=0.0262, p=0.9795). E) Rectal body temperature at baseline, during SEFL, and post-SEFL, in Ctrl-SEFL and Cre-SEFL animals (n=7-9, two-way ANOVA, F(2,41)=0.7207, p=0.4925). F–G) Quantification of fat mass (F) (n=12-14, unpaired t-test, t=0.397, p=0.7080) and lean mass (G) (n=12-14, unpaired t-test, t=0.6018, p=0.5529) in Ctrl-SEFL and Cre-SEFL groups. H–I) Food consumption (H) and locomotor activity (I) across full-day, dark, and light phases in all four experimental groups (Ctrl-No SEFL, Ctrl-SEFL, Cre-No SEFL, Cre-SEFL) (n=7-10, one-way ANOVA, p>0.05). J) Oxygen consumption (VO₂) (n=5, p=0.6191) and energy expenditure (EE) (n=5, p=0.661) under cold conditions (4^0^C) in Ctrl-SEFL and Cre-SEFL groups.K) Representative images of liver tissue before and after clearing, with corresponding 3D reconstructions of TH immunolabeling in Ctrl-SEFL and Cre-SEFL groups. L) Higher-magnification cleared liver images showing TH expression.M) Quantification of normalized TH intensity in liver tissue sample of Ctrl-SEFL and Cre-SEFL groups (n=2, unpairede t-test, t=0.01698, p=0.9865).N) Pyruvate tolerance test measuring blood glucose over time in Ctrl-SEFL and Cre-SEFL groups (n=7-8, two-way ANOVA, F(1,78)=0.2053, p=0.6517).O) Plasma insulin levels measured at baseline and following glucose injection at 10 and 30 minutes (n=6-8, two-way ANOVA, F(1,37)=1.403, p=0.2347).P) Area under the curve (AUC) quantification of insulin response (n=6-8, unpaired t-test, t=0.1170, p=0.9088).

## References

Abreu-Vieira, G., Xiao, C., Gavrilova, O., and Reitman, M.L. (2015). Integration of body temperature into the analysis of energy expenditure in the mouse. Mol Metab 4, 461–470.

Andrews, R.C., Herlihy, O., Livingstone, D.E., Andrew, R., and Walker, B.R. (2002). Abnormal cortisol metabolism and tissue sensitivity to cortisol in patients with glucose intolerance. J Clin Endocrinol Metab 87, 5587–5593.

Andrikopoulos, S., Blair, A.R., Deluca, N., Fam, B.C., and Proietto, J. (2008). Evaluating the glucose tolerance test in mice. Am J Physiol Endocrinol Metab 295, E1323–1332.

Bamshad, M., Song, C.K., and Bartness, T.J. (1999). CNS origins of the sympathetic nervous system outflow to brown adipose tissue. Am J Physiol 276, R1569–1578.

Bartness, T.J., Vaughan, C.H., and Song, C.K. (2010). Sympathetic and sensory innervation of brown adipose tissue. Int J Obes (Lond) 34 *Suppl 1*, S36–42.

Benjamini, Y., and Hochberg, Y. (1995). Controlling the False Discovery Rate: A Practical and Powerful Approach to Multiple Testing. Journal of the Royal Statistical Society Series B (Methodological) 57, 289–300.

Blazer, D.G., and Hybels, C.F. (2010). Shortness of breath as a predictor of depressive symptoms in a community sample of older adults. Int J Geriatr Psychiatry 25, 1080–1084.

Blechert, J., Michael, T., Grossman, P., Lajtman, M., and Wilhelm, F.H. (2007). Autonomic and respiratory characteristics of posttraumatic stress disorder and panic disorder. Psychosom Med 69, 935–943.

Blenkuš, U., Gerós, A.F., Carpinteiro, C., Aguiar, P.C., Olsson, I.A.S., and Franco, N.H. (2022). Non-Invasive Assessment of Mild Stress-Induced Hyperthermia by Infrared Thermography in Laboratory Mice. Animals (Basel) 12.

Borgdorff, P. (1975). Respiratory fluctuations in pupil size. Am J Physiol 228, 1094–1102.

Bozadjieva-Kramer, N., Ross, R.A., Johnson, D.Q., Fenselau, H., Haggerty, D.L., Atwood, B., Lowell, B., and Flak, J.N. (2021). The Role of Mediobasal Hypothalamic PACAP in the Control of Body Weight and Metabolism. Endocrinology 162.

Brown, R.P., and Gerbarg, P.L. (2005). Sudarshan Kriya Yogic breathing in the treatment of stress, anxiety, and depression. Part II--clinical applications and guidelines. J Altern Complement Med 11, 711–717.

Brown, R.P., Gerbarg, P.L., and Muench, F. (2013). Breathing practices for treatment of psychiatric and stress-related medical conditions. Psychiatr Clin North Am 36, 121–140.

Clancy, K.J., Devignes, Q., Kumar, P., May, V., Hammack, S.E., Akman, E., Casteen, E.J., Pernia, C.D., Jobson, S.A., Lewis, M.W., et al. (2023). Circulating PACAP levels are associated with increased amygdala-default mode network resting-state connectivity in posttraumatic stress disorder. Neuropsychopharmacology 48, 1245–1254.

de Sousa Abreu, R.P., Hoffman, A.N., Bondarenko, E., Huang, Y., Burgos Pujols, R.E., Fanselow, M.S., and Feldman, J.L. (2024). Episodic slow breathing in mice markedly reduces fear responses. bioRxiv, 2024.2012.2009.627565.

Dominelli, P.B., and Sheel, A.W. (2024). The pulmonary physiology of exercise. Adv Physiol Educ 48, 238–251.

Duesman, S.J., Shetty, S., Patel, S., Ogale, N., Mohamed, F., Sparman, N., Rajbhandari, P., and Rajbhandari, A.K. (2022). Sexually dimorphic role of the locus coeruleus PAC1 receptors in regulating acute stress-associated energy metabolism. Frontiers in Behavioral Neuroscience 16.

Fehr, F.S., and Stern, J.A. (1970). Peripheral physiological variables and emotion: the James-Lange theory revisited. Psychol Bull 74, 411–424.

Feldman, J.L., and Del Negro, C.A. (2006). Looking for inspiration: new perspectives on respiratory rhythm. Nat Rev Neurosci 7, 232–242.

Feldman, J.L., Del Negro, C.A., and Gray, P.A. (2013). Understanding the rhythm of breathing: so near, yet so far. Annu Rev Physiol 75, 423–452.

Feldman, J.L., and Smith, J.C. (1989). Cellular mechanisms underlying modulation of breathing pattern in mammals. Ann N Y Acad Sci 563, 114–130.

Fichna, M., and Fichna, P. (2017). Glucocorticoids and beta-cell function. Endokrynol Pol 68, 568–573.

First, M.B. (2013). Diagnostic and statistical manual of mental disorders, 5th edition, and clinical utility. J Nerv Ment Dis 201, 727–729.

Garfinkel, S.N., and Critchley, H.D. (2016). Threat and the Body: How the Heart Supports Fear Processing. Trends Cogn Sci 20, 34–46.

Garfinkel, S.N., Minati, L., Gray, M.A., Seth, A.K., Dolan, R.J., and Critchley, H.D. (2014). Fear from the heart: sensitivity to fear stimuli depends on individual heartbeats. J Neurosci 34, 6573–6582.

Gray, P.A., Janczewski, W.A., Mellen, N., McCrimmon, D.R., and Feldman, J.L. (2001). Normal breathing requires preBötzinger complex neurokinin-1 receptor-expressing neurons. Nat Neurosci 4, 927–930.

Hahn, M.K., Giacca, A., and Pereira, S. (2024). In vivo techniques for assessment of insulin sensitivity and glucose metabolism. J Endocrinol 260.

Hamasaki, H. (2020). Effects of Diaphragmatic Breathing on Health: A Narrative Review. Medicines (Basel) 7.

Hammack, S.E., Cheung, J., Rhodes, K.M., Schutz, K.C., Falls, W.A., Braas, K.M., and May, V. (2009). Chronic stress increases pituitary adenylate cyclase-activating peptide (PACAP) and brain-derived neurotrophic factor (BDNF) mRNA expression in the bed nucleus of the stria terminalis (BNST): roles for PACAP in anxiety-like behavior. Psychoneuroendocrinology 34, 833–843.

Hao, Y., Stuart, T., Kowalski, M.H., Choudhary, S., Hoffman, P., Hartman, A., Srivastava, A., Molla, G., Madad, S., Fernandez-Granda, C., et al. (2024). Dictionary learning for integrative, multimodal and scalable single-cell analysis. Nat Biotechnol 42, 293–304.

Heck, D.H., Kozma, R., and Kay, L.M. (2019). The rhythm of memory: how breathing shapes memory function. J Neurophysiol 122, 563–571.

Hsueh, B., Chen, R., Jo, Y., Tang, D., Raffiee, M., Kim, Y.S., Inoue, M., Randles, S., Ramakrishnan, C., Patel, S., et al. (2023). Cardiogenic control of affective behavioural state. Nature 615, 292–299.

James-Lange (1922). The emotions, Vol. 1 (Baltimore, MD, US: Williams & Wilkins Co).

Klein, A.S., Dolensek, N., Weiand, C., and Gogolla, N. (2021). Fear balance is maintained by bodily feedback to the insular cortex in mice. Science 374, 1010–1015.

Klein, D.F. (1993). False suffocation alarms, spontaneous panics, and related conditions. An integrative hypothesis. Arch Gen Psychiatry 50, 306–317.

Kosmidis, S., Negrean, A., Dranovsky, A., Losonczy, A., and Kandel, E.R. (2021). A fast, aqueous, reversible three-day tissue clearing method for adult and embryonic mouse brain and whole body. Cell Rep Methods 1, 100090.

Kuo, T., McQueen, A., Chen, T.C., and Wang, J.C. (2015). Regulation of Glucose Homeostasis by Glucocorticoids. Adv Exp Med Biol 872, 99–126.

Ley, R. (1988). Panic attacks during relaxation and relaxation-induced anxiety: a hyperventilation interpretation. J Behav Ther Exp Psychiatry 19, 253–259.

Litz, B.T., Weathers, F.W., Monaco, V., Herman, D.S., Wulfsohn, M., Marx, B., and Keane, T.M. (1996). Attention, arousal, and memory in posttraumatic stress disorder. J Trauma Stress 9, 497–519.

Melnychuk, M.C., Dockree, P.M., O’Connell, R.G., Murphy, P.R., Balsters, J.H., and Robertson, I.H. (2018). Coupling of respiration and attention via the locus coeruleus: Effects of meditation and pranayama. Psychophysiology 55, e13091.

Michopoulos, V., Vester, A., and Neigh, G. (2016). Posttraumatic stress disorder: A metabolic disorder in disguise? Exp Neurol 284, 220–229.

Mina, A.I., LeClair, R.A., LeClair, K.B., Cohen, D.E., Lantier, L., and Banks, A.S. (2018). CalR: A Web-Based Analysis Tool for Indirect Calorimetry Experiments. Cell Metab 28, 656–666.e651.

Mustafa, T. (2013). Pituitary adenylate cyclase-activating polypeptide (PACAP): a master regulator in central and peripheral stress responses. Adv Pharmacol 68, 445–457.

Nagai, J., Rajbhandari, A.K., Gangwani, M.R., Hachisuka, A., Coppola, G., Masmanidis, S.C., Fanselow, M.S., and Khakh, B.S. (2019). Hyperactivity with Disrupted Attention by Activation of an Astrocyte Synaptogenic Cue. Cell 177, 1280–1292.e1220.

Nishimura, K.J., Poulos, A.M., Drew, M.R., and Rajbhandari, A.K. (2022). Know thy SEFL: Fear sensitization and its relevance to stressor-related disorders. Neurosci Biobehav Rev 142, 104884.

Perusini, J.N., Meyer, E.M., Long, V.A., Rau, V., Nocera, N., Avershal, J., Maksymetz, J., Spigelman, I., and Fanselow, M.S. (2016). Induction and Expression of Fear Sensitization Caused by Acute Traumatic Stress. Neuropsychopharmacology 41, 45–57.

Poulos, A.M., Zhuravka, I., Long, V., Gannam, C., and Fanselow, M. (2015). Sensitization of fear learning to mild unconditional stimuli in male and female rats. Behav Neurosci 129, 62–67.

Rajbhandari, A.K., Gonzalez, S.T., and Fanselow, M.S. (2018). Stress-Enhanced Fear Learning, a Robust Rodent Model of Post-Traumatic Stress Disorder. J Vis Exp.

Rajbhandari, A.K., Octeau, C.J., Gonzalez, S., Pennington, Z.T., Mohamed, F., Trott, J., Chavez, J., Ngyuen, E., Keces, N., Hong, W.Z., et al. (2021). A Basomedial Amygdala to Intercalated Cells Microcircuit Expressing PACAP and Its Receptor PAC1 Regulates Contextual Fear. J Neurosci 41, 3446–3461.

Rau, V., DeCola, J.P., and Fanselow, M.S. (2005). Stress-induced enhancement of fear learning: an animal model of posttraumatic stress disorder. Neurosci Biobehav Rev 29, 1207–1223.

Ressler, K.J., Mercer, K.B., Bradley, B., Jovanovic, T., Mahan, A., Kerley, K., Norrholm, S.D., Kilaru, V., Smith, A.K., Myers, A.J., et al. (2011). Post-traumatic stress disorder is associated with PACAP and the PAC1 receptor. Nature 470, 492–497.

Rosenbaum, S., Stubbs, B., Ward, P.B., Steel, Z., Lederman, O., and Vancampfort, D. (2015). The prevalence and risk of metabolic syndrome and its components among people with posttraumatic stress disorder: a systematic review and meta-analysis. Metabolism 64, 926–933.

Schwarzacher, S.W., Rüb, U., and Deller, T. (2011). Neuroanatomical characteristics of the human pre-Bötzinger complex and its involvement in neurodegenerative brainstem diseases. Brain 134, 24–35.

Shepherd, L., and Wild, J. (2014). Emotion regulation, physiological arousal and PTSD symptoms in trauma-exposed individuals. J Behav Ther Exp Psychiatry 45, 360–367.

Shi, Y., Stornetta, D.S., Reklow, R.J., Sahu, A., Wabara, Y., Nguyen, A., Li, K., Zhang, Y., Perez-Reyes, E., Ross, R.A., et al. (2021). A brainstem peptide system activated at birth protects postnatal breathing. Nature 589, 426–430.

Smith, J.C., Ellenberger, H.H., Ballanyi, K., Richter, D.W., and Feldman, J.L. (1991). Pre-Bötzinger complex: a brainstem region that may generate respiratory rhythm in mammals. Science 254, 726–729.

Tan, W., Pagliardini, S., Yang, P., Janczewski, W.A., and Feldman, J.L. (2010). Projections of preBötzinger complex neurons in adult rats. J Comp Neurol 518, 1862–1878.

Tjondrorahardja, E.J., Poon, T.T.S., Avraam, J., Schenker, M., Felmingham, K.L., and Jordan, A.S. (2024). Breathlessness, but not breathing instability, varies with posttraumatic stress symptoms in university students. J Appl Physiol (1985) 137, 1458–1469.

Tong, A.J., Liu, X., Thomas, B.J., Lissner, M.M., Baker, M.R., Senagolage, M.D., Allred, A.L., Barish, G.D., and Smale, S.T. (2016). A Stringent Systems Approach Uncovers Gene-Specific Mechanisms Regulating Inflammation. Cell 165, 165–179.

Van der Heyden, J.A., Zethof, T.J., and Olivier, B. (1997). Stress-induced hyperthermia in singly housed mice. Physiol Behav 62, 463–470.

van Liempt, S., Westenberg, H.G., Arends, J., and Vermetten, E. (2011). Obstructive sleep apnea in combat-related posttraumatic stress disorder: a controlled polysomnography study. Eur J Psychotraumatol 2.

Virtue, S., and Vidal-Puig, A. (2021). GTTs and ITTs in mice: simple tests, complex answers. Nat Metab 3, 883–886.

von Leupoldt, A., Chan, P.Y., Bradley, M.M., Lang, P.J., and Davenport, P.W. (2011). The impact of anxiety on the neural processing of respiratory sensations. Neuroimage 55, 247–252.

Von Leupoldt, A., Vovk, A., Bradley, M.M., Keil, A., Lang, P.J., and Davenport, P.W. (2010). The impact of emotion on respiratory-related evoked potentials. Psychophysiology 47, 579–586.

Williams, L.M., Campbell, F.M., Drew, J.E., Koch, C., Hoggard, N., Rees, W.D., Kamolrat, T., Thi Ngo, H., Steffensen, I.L., Gray, S.R., et al. (2014). The development of diet-induced obesity and glucose intolerance in C57BL/6 mice on a high-fat diet consists of distinct phases. PLoS One 9, e106159.

Yang, C.F., and Feldman, J.L. (2018). Efferent projections of excitatory and inhibitory preBötzinger Complex neurons. J Comp Neurol 526, 1389–1402.

Yang, C.F., Kim, E.J., Callaway, E.M., and Feldman, J.L. (2020). Monosynaptic Projections to Excitatory and Inhibitory preBötzinger Complex Neurons. Front Neuroanat 14, 58.

Yehuda, R. (2002). Post-Traumatic Stress Disorder. New England Journal of Medicine 346, 108–114.

Yoshimoto, A., Morikawa, S., Kato, E., Takeuchi, H., and Ikegaya, Y. (2024). Top-down brain circuits for operant bradycardia. Science 384, 1361–1368.

Zelano, C., Jiang, H., Zhou, G., Arora, N., Schuele, S., Rosenow, J., and Gottfried, J.A. (2016). Nasal Respiration Entrains Human Limbic Oscillations and Modulates Cognitive Function. J Neurosci 36, 12448–12467.

